# Sensitive Spatiotemporal Tracking of Spontaneous Metastasis in Deep Tissues via a Genetically-Encoded Magnetic Resonance Imaging Reporter

**DOI:** 10.1101/2022.02.09.479610

**Authors:** Nivin N. Nyström, Sean W. McRae, Francisco F.M. Martinez, John J. Kelly, Timothy J. Scholl, John A. Ronald

**Author notes:** Correspondence can be addressed to J.A.R. and T.J.S.

## Abstract

Metastasis remains a poorly understood aspect of cancer biology and the leading cause of cancer-related death, yet most preclinical cancer studies do not examine metastasis, focusing solely on the primary tumor. One major factor contributing to this paradox is a gap in available tools for accurate spatiotemporal measurements of metastatic spread *in vivo*. Our objective was to develop an imaging reporter system that offers sensitive three-dimensional detection of cancer cells at high resolutions in live mice. We utilized *organic anion-transporting polypeptide lb3* (*oatp1b3*) as a magnetic resonance imaging (MRI) reporter gene to this end, and systematically optimized its framework for *in vivo* tracking of viable cancer cells in a spontaneous metastasis model. We were able to image metastasis on *oatp1b3*-MRI at the single lymph node level and continued to track its progression over time as cancer cells spread to multiple lymph nodes and different organ systems in single animals. While initial single lesions were successfully imaged in parallel via bioluminescence, later metastases were obscured by light scatter from the initial node. Importantly, we demonstrate and validate that 100-μm isotropic resolution MR images could detect micrometastases in lung tissue estimated to contain fewer than 10^3^ cancer cells. In summary, *oatp1b3*-MRI enables precise determination of lesion size and location over time and offers a path towards deep-tissue tracking of any *oatp1b3-engineered* cell type with combined high resolution, high sensitivity, 3D spatial information, and surrounding anatomical context.

## INTRODUCTION

Metastasis is responsible for approximately 90% of cancer-related mortalities, yet it remains the least understood aspect of cancer biology^1^. Preclinical animal studies provide a valuable platform for investigating metastasis, intermediate to reductionist *in vitro* models and expensive clinical trials^2^. Still, approximately 75% of preclinical animal studies published in leading cancer journals do not investigate metastasis to any extent, instead focusing only on the primary tumor, largely due to a lack of methods for accurate spatiotemporal quantification of metastatic burden^3^. Spontaneous metastasis models, which recapitulate the entire metastatic cascade and better mimic clinical disease, are even rarer in the literature, as they complicate experiments further with increased variability between animals in both the rate and site pattern of metastatic progression^4^.

Bioluminescence imaging (BLI) is routinely used for assessing whole animal burden in experimental metastasis models because of its high throughput and sensitivity; but assessing total burden on BLI with accuracy, especially in spontaneous metastasis models that include primary tumors, remains a challenge. Light scatter from larger lesions and light attenuation by surrounding tissues contribute to poor resolution, signal loss and/or obscurement of smaller or more deep-seated metastases^5^. With current imaging methods, lesion size, depth and precise location remain unclear prior to posthumous examination. Preclinical studies have therefore paired BLI with tissue clearing protocols, light-sheet microscopy and deep-learning methods for sensitive imaging of metastatic cells at endpoint^6–9^. Alternate to BLI, other approaches include implantation of permanent optical windows for real-time monitoring of specific tissue sites^10^ or reverting to non-mammalian model organisms to accomplish longitudinal, high resolution, *in vivo* imaging of metastasis^11^. All the while, sensitive, high resolution *in vivo* tracking of metastasis in deep tissues remains an elusive goal^2^.

Magnetic resonance imaging (MRI) uniquely provides high resolution, 3-dimentional spatial information with excellent soft tissue contrast, and is extensively used for preclinical assessment of primary tumors and metastatic lesions^12,13^. Although MRI offers versatile contrast mechanisms to enhance lesions on images, it still faces challenges in detecting small metastases due to its relatively low sensitivity; state-of-the-art MR imaging probes require that lesions grow to diameters of at least ~0.5 mm in lungs of mice, and require even larger diameters at other tissue sites^14,15^. Reporter genes for MRI have previously been developed to enhance contrast of cells of interest^16–22^, but have not been seriously explored for preclinical tracking of metastasis.

For example, *organic anion-transporting polypeptide 1* (*oatp1*), which encodes a 12-transmembrane-domain integral membrane protein, was previously established as a reporter gene based on its ability to bind and transport gadolinium ethoxybenzyl diethylene triamine pentaacetic acid (Gd-EOB-DTPA) from the extracellular environment into *oatp1*-engineered cells^23^. We and others applied *oatp1*-MRI to *in vivo* cancer imaging and showcased its ability to generate detailed, high-resolution images of primary tumor architecture in mice^24^, but measured its *in vivo* detection limit at 10^6^ cells per lesion^25^. Meanwhile, non-MR reporter genes recently achieved detection of single isolated cells on BLI^26^ and detection of lesions on the order of 10^4^ cells on positron emission tomography (PET)^27^.

However, with multiple variables contributing to *oatp1*-MRI performance, from genetic construct design to biophysical imaging parameters, we hypothesized that systematic optimization of the imaging framework could greatly improve its *in vivo* detection limit. Accordingly, our primary objective was to increase the sensitivity of the *oatp1*-MRI system for detection and dynamic tracking of *oatp1*-engineered cancer cells in a spontaneous metastasis model of breast cancer. We report that *oatp1*-MRI produces highly-sensitive, 3-dimensional, and high-resolution images of metastatic progression in live mice over time; we improve system sensitivity by three orders of magnitude relative to previous studies^23–25^, enabling detection of fewer than 10^3^ cells per lesion, or 10^2^ cells per voxel, in lung tissue. Importantly, *oatp1*-MRI enables detection of reporter gene signals that are unaffected by tissue depth or the presence of adjacent lesions. The information afforded by *oatp1*-MRI thus enables precise determination of lesion size, depth and location for spatiotemporal profiling of metastatic burden in deep tissues of live animals.

## RESULTS

### *In silico* and *in vitro* protein characterization of OATP1B3 synthetically expressed in a metastasis-competent cancer cell line

Two lentiviral transfer plasmids, co-encoding *tdTomato fluorescent protein* with *luciferase* and *zsGreen fluorescent protein* with *oatp1b3* were cloned and packaged into lentiviral vectors (**Figure 1b**). Both transgene cassettes were placed under regulation of the constitutive human elongation factor-1 alpha promoter (pEF1α) to maximize expression. A self-cleaving peptide (P2A, T2A, respectively) was used to incorporate fluorescent proteins for fluorescence-activated cell sorting (FACS) and a woodchuck hepatitis virus post-transcriptional regulatory element (WPRE) was used to stabilize mRNA transcript levels. Metastasis-competent human triple negative breast cancer cells (MDA-MB-231) were transduced with fresh lentivirus and subsequently sorted with gates for high expression (top 3%) of both tdTomato and zsGreen fluorescence with >98% purity to obtain stable *luciferase-expressing* control cells (Luc-CTL), as well as cells with stable co-expression of *luciferase* and *oatp1b3* (Luc-1B3).

**Figure 1.**
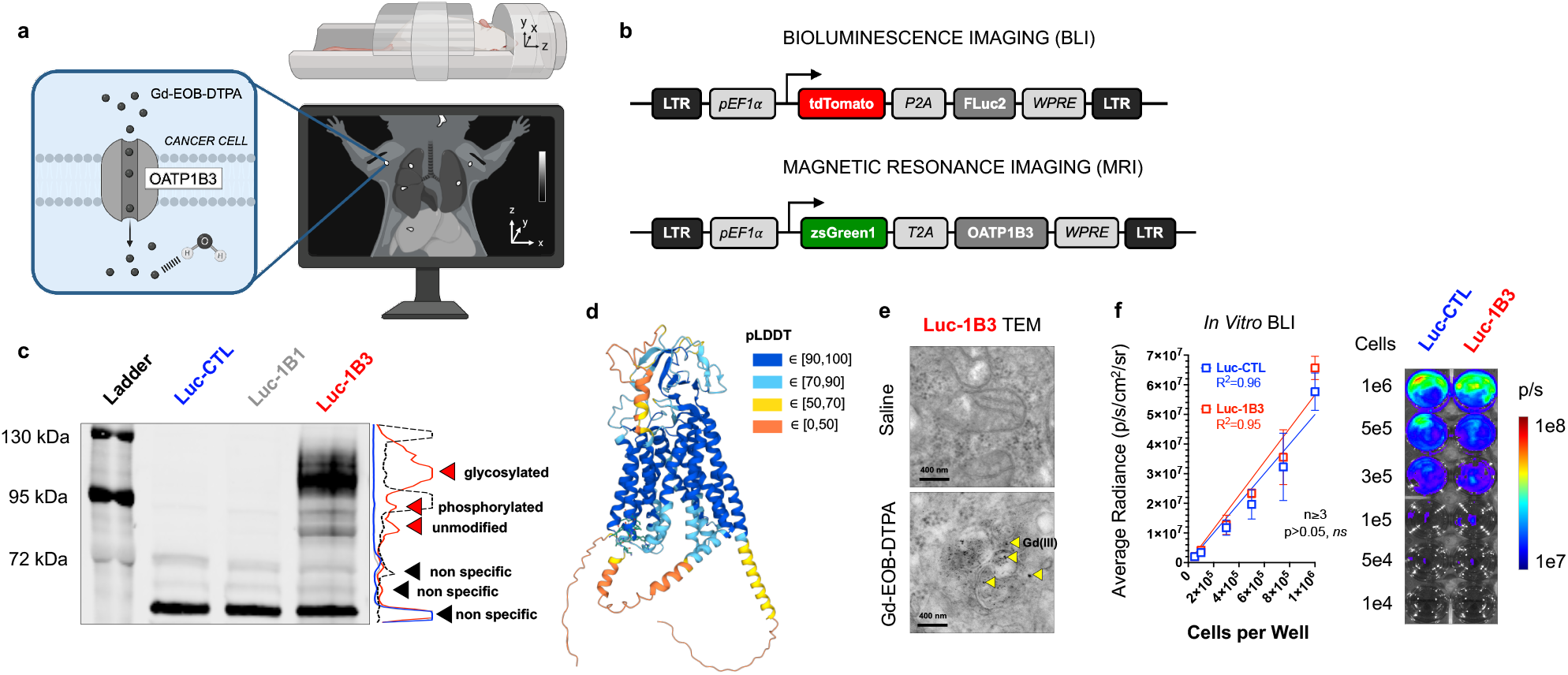
Principle of *Oatp1b3* as a Reporter Gene for Cancer Cell Tracking. a) Synthetic expression of the OATP1B3 transporter by cancer cells enables Gd-EOB-DTPA uptake into the cellular cytoplasm compartment of cells. This causes increased spin-lattice relaxation of water protons, which can be detected with *T*_1_-weighted magnetic resonance imaging producing high-resolution, 3-dimensional images with anatomical context. b) Genetic constructs for transgene expression via lentiviral integration. For bioluminescence imaging (BLI), human codon-optimized firefly luciferase (FLuc2) was co-encoded with tdTomato fluorescent protein, which was used as a marker for cell sorting. For magnetic resonance imaging, *Organic anion-transporting polypeptide lb3* (OATP1B3) was co-encoded with zsGreen1 fluorescent protein for cell sorting, long terminal repeat, LTR, human elongation factor-1 alpha promoter, pEF1α, self-cleaving peptides, P2A, T2A, and woodchuck hepatitis virus post-transcriptional regulatory element, WPRE. c) Anti-OATP1B3 Western blot of cells engineered with luciferase (Luc-CTL, *blue)*, cells engineered with luciferase and *oatp1bl* (Luc-1B1, *gray*), and cells engineered with luciferase and *oatp1b3* (Luc-1B3, *red*). Signal intensity profiles are outlined along position of Western blot. Ladder profile is shown in black. Non-specific peaks and OATP1B3-specific peaks of unmodified, phosphorylated, and glycosylated OATP1B3 are indicated by arrows. d) 3D rendering of AlphaFold^32^ prediction for OATP1B3 structure, illustrated with single residue-resolved confidence score. pLDDT, predicted local-distance difference test. e) Transmission electron microscopy of Luc-1B3 cells incubated with or without 1 mM Gd-EOB-DTPA for 1 hour. Gd(III) appears as black foci, indicated by yellow arrows. f) Average radiance (p/s/cm^2^/sr) of increasing numbers of Luc-CTL (*blue*) and Luc-1B3 cells (*red*) per well, n=3, p>0.05, ns. Representative image of well plate. Error bars represent one standard deviation.

Immuno-blotting of cell lysates for OATP1B3 protein confirmed its absence in Luc-CTL cells, whereas four peaks at distinct molecular weights, measured at 84 kDa, 92 kDa, 112 kDa, and 120 kDa developed in the Luc-1B3 lane (**Figure 1c**). Non-specific bands across all samples were observed at about 60 kDa, 65 kDa and 72 kDa (**Figure 1c**). Some post-translational modifications of OATP1B3 have previously been reported; namely, two glycosylation sites have been associated with its trafficking to the cell membrane and its functionalization^28^, whereas increased phosphorylation of OATP1B3 was correlated with downregulation of its transport activity^29^. Band analysis with FindMod-ExPASy^30^ (expasy.org, Swiss Institute of Bioinformatics, Lausanne Switzerland) resulted in identification of the 84 kDa band as the unmodified transporter, the band at about 92 kDa as the phosphorylated protein, and the wide band about 112 kDa as the glycosylated transporter with >95% confidence. Finally, the less intense peak at about 120 kDa was identified as OATP1B3 that is both glycosylated and phosphorylated, albeit with <95% confidence^31^. Importantly, the Western blot suggests that a significant proportion of the protein synthesized by engineered MDA-MB-231 cells has undergone the glycosylation necessary for functionalization (75.8%, AUC = 40.9 a.u.), whereas a smaller fraction (12.3%, AUC = 6.61 a.u.) was phosphorylated but not glycosylated (**Figure 1c**).

AlphaFold^32^ was employed to predict molecular structure of OATP1B3 (Identifier AF-Q9NPD5-F1), generating a model with an overall confidence score (predicted local distance difference test, pLDDT) of 77.6% (**Figure 1d, Supplementary Figure S1**). Processing of the OATP1B3 protein sequence and structure (UniProtKB ID SO1B3_HUMAN, UniProtKB Accession Q9NPD5-1), first through iPTMnet^33^ (research.bioinformatics.udel.edu/iptmnet/, University of Delaware, Newark, DE), then subsequently via GlyGen^34^ (glygen.org, University of Georgia, Athens, GA) and PhosphoSitePlus^35^ (phosphosite.org, Cell Signaling Technology, Danvers, MA) resulted in identification of 8 extracellular N-linked glycosylation sites and 3 intracellular phosphoserine sites, as well as an intracellular serine protease inhibitor motif characterized by 3 disulfide bonds (**Table S1**).

Transmission electron microscopy of Luc-1B3 cells incubated with Gd-EOB-DTPA confirmed the influx of Gd(III) into the cytoplasmic space of Luc-1B3 cells, where the paramagnetic center would be free to interact with protons of the intracellular environment (**Figure 1e**). Curiously, Gd(III) was also found encapsulated within residual bodies, destined for exocytosis, suggesting a distinct elimination pathway that to our knowledge has not previously been reported (**Supplementary Figure S2**). Finally, BLI demonstrated a strong positive linear correlation between cell number and average radiance (p/s/cm^2^/sr) for both Luc-CTL (R^2^=0.96) and Luc-1B3 (R^2^=0.95) cells *in vitro*. The slope of the linear regression was not significantly different between Luc-CTL (55.2 ±8.1 p/s/cm^2^/sr/cell) and Luc-1B3 cells (62.5 ± 1.9 p/s/cm^2^/sr/cell) (p=0.22, **Figure 1f**). This serves as an important control for BLI in later animal experiments.

### Solving for optimal parameters of imaging for *oatp1b3-MRI*

After cell engineering, we first set out to characterize the biochemical, kinetic and magnetic relaxation properties of the *oatp1b3* reporter gene system *in vitro* (**Figure 2**). First, Gd-EOB-DTPA uptake in Luc-CTL and Luc-1B3 cells was measured as a function of applied concentration via inductively coupled plasma mass spectrometry (**Figure 2a**). A significant increase in intracellular Gd(III) was observed in Luc-1B3 cells at all applied concentrations relative to Luc-CTL cells (p<0.05, n=3), but no significant difference was observed between Luc-CTL cells treated with increasing Gd-EOB-DTPA concentrations (p>0.05, n=3), suggesting that Gd-EOB-DTPA is highly specific to *oatp1b3* and its uptake by non *oatp1b3*-expressing cells is negligible within this timeframe (**Figure 2a**). After fitting the data to an exponential plateau model (R^2^=0.81), the maximum intracellular capacity of Gd(III) for our system was calculated to be 1.66 μg Gd(III) per million cells, or approximately 1.66 pg Gd(III) per single cell (**Figure 2a**), which signified that channel-mediated transport of Gd-EOB-DTPA into cells constitutively expressing *oatp1b3* approaches saturation (99.6 ± 18.0% of maximum capacity) at 16 mM Gd-EOB-DTPA.

**Figure 2.**
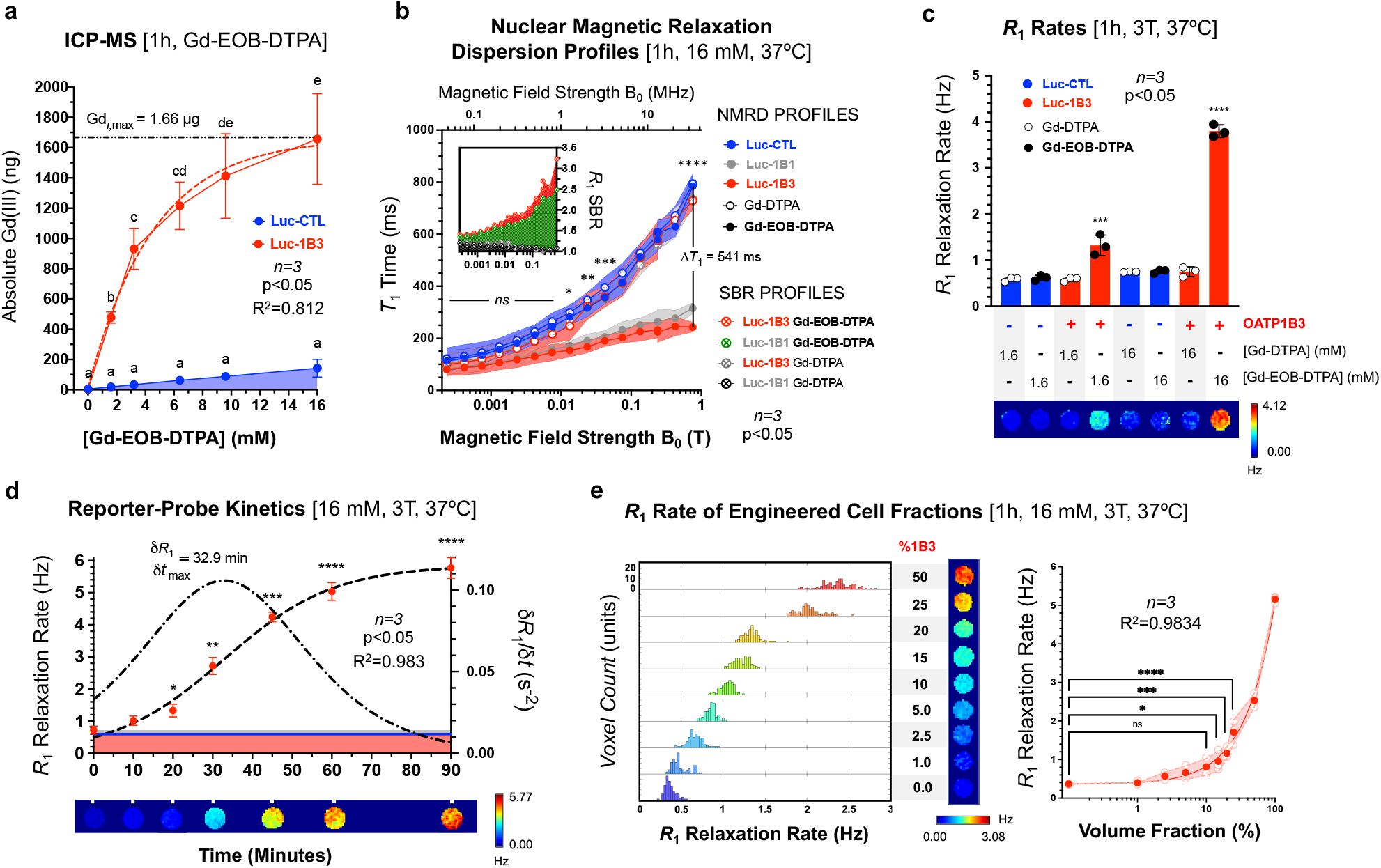
Characterization of Biochemical, Magnetic Relaxation and Kinetic Properties of Gd-EOB-DTPA in a System of Mammalian Cells Expressing *Oatp1b3*. a) Inductively-coupled plasma mass spectrometry (ICP-MS) of Gd(III) (ng) from lysate of a million Luc-CTL (*blue*) and Luc-1B3 cells (*red*) incubated with variable [Gd-EOB-DTPA] for 1 hour, n=3, *p<0.05, R^2^=0.82. Gd_i,max_, maximum intracellular Gd(III) mass projected (per million cells). b) *T*_1_ time (ms) measured at variable field strength (MHz) of Luc-CTL (*blue)*, Luc-1B1 (*grey)*, and Luc-1B3 cells (*red*) incubated with either 16 mM Gd-DTPA 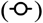 or Gd-EOB-DTPA 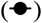 for 1 hour. Signal-to-background ratio (SBR; arbitrary units, a.u.) of *R*_1_ relaxation rates of Luc-1B1 (*green*) and Luc-1B3 (*red*) cells incubated with Gd-EOB-DTPA are shown in the embedded graph. Background is Luc-CTL cells at the same field strength and probe condition. n=3, *p<0.05. c) relaxation rates (Hz) measured at 3T and 37°C of Luc-CTL (*blue*) and Luc-1B3 cells (*red*) incubated with Gd-DTPA 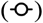 or Gd-EOB-DTPA 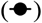 for 1 hour. n=3, *p<0.05. d) relaxation rates (Hz) of Luc-1B3 cells at 3T incubated with 16 mM Gd-EOB-DTPA for variable incubation times at 37°C. n=3, *p<0.05. Regression of Luc-1B3 *R*_1_ rates 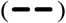, the first derivative of the regression 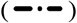, Luc-CTL cells incubated with Gd-EOB-DTPA for 90 minutes (*blue)*, Luc-1B3 cells incubated with Gd-DTPA for 90 minutes (*pink)*, and phosphate buffered saline (*grey*) are also plotted. e) relaxation rates (Hz) of variable Luc-CTL and Luc-1B3 cell ratios at and 37°C incubated with 16 mM Gd-EOB-DTPA for 1 hour. Filled circles (*red*) represent average values, open circles (*pink*) are individual trial measurements. Shading represents standard deviation. n=3, *p<0.05, R^2^=0.98.

Although the relaxivity of Gd(III)-based contrast agents, including Gd-EOB-DTPA, decreases as a function of field strength and therefore exhibits its greatest effects at low field (B_0_ < 0.05 T; **Supplementary Figure S3**), tissue *R*_1_ values also decrease along the same axis, and these competing phenomena both affect the resultant contrast enhancement that is central to our endeavor of detecting metastasis *in vivo* with high sensitivity. As expected of non-engineered tissues, with increasing field strength, where B_0_ ∈ (0.000233, 1.0009) T, we observed significant increases in the *T*_1_ times of all cells treated with the control Gd-DTPA probe (7.29-fold, p<0.0001) and Luc-CTL cells treated with the *oatp1b3*-targeted Gd-EOB-DTPA (6.94-fold, p<0.0001; **Figure 2b**). For Luc-1B3 cells incubated with Gd-EOB-DTPA, Gd(III) becomes a major contributing factor to spin-lattice relaxation time; the *T*_1_ time still increased as a function of field strength (3.04-fold), but was significantly lower than that of control samples (n=3, p<0.0001; **Figure 2b**). Notably, the difference in *T*_1_ time between Luc-CTL and Luc-1B3 cells both treated with Gd-EOB-DTPA significantly increased with magnetic field strength (n=3, p<0.05; **Figure 2b**). At the lower field limit of 0.000233 T, △*T*_1_ between Luc-CTL and Luc-1B3 cells was 33.1 ± 4.0 ms but this difference increased to 541 ± 20 ms at the upper field limit of 1.0009 T (n=3, p<0.0001; **Figure 2b**).

At a field strength of 3 Tesla, Luc-1B3 cells incubated with low (1.6 mM) and high (16 mM) Gd-EOB-DTPA concentrations exhibited significantly increased *R*_1_ rates (1.32 ± 0.22, 3.80 ± 0.14 s^-1^, respectively) compared to all other control conditions (n=3, p≤0.0001; **Figure 2c**). Specifically, Luc-1B3 cells exhibited a 2.14 *R*_1_ signal-to-background ratio (SBR) at 1.6 mM Gd-EOB-DTPA concentrations and a more substantial 5.02-fold *R*_1_ SBR at 16 mM Gd-EOB-DTPA concentrations (**Figure 2c)**, which was also greater than the SBR measured at lower fields (SBR_1T_ = 3.23; **Figure 2b**). At fields greater than 3T, however, Gd-EOB-DTPA relaxivity continues to decrease (**Supplementary Figure S3**) while spin-lattice relaxation rates for all tissues become similar as they approach zero, resulting in the marked reduction of contrast enhancement typically observed in high field MRI (*i.e*. 7T, 9.4T)^36,37^. We therefore reasoned that imaging animals at mid-field (*i.e*. 3T) would optimize SBR for *in vivo* detection of metastases. All the while, to maximize detection sensitivity while mitigating any toxicity concerns from using concentrations of Gd-EOB-DTPA higher than 16 mM, which would result in only incremental increases to SBR (**Figure 2a**), we selected an applied concentration of 16 mM Gd-EOB-DTPA for all *in vitro* MRI experiments, and a correspondent dose of 1.3 mmol/kg Gd-EOB-DTPA for *in vivo* MRI experiments.

When Luc-1B3 cells were incubated with Gd-EOB-DTPA for variable lengths of time, significant increases in *R*_1_ relative to Luc-CTL cells were first observed at the 20-minute timepoint (1.33 ± 0.20 Hz; n=3, p=0.0015; **Figure 2d**). The *R*_1_ of Luc-1B3 cells as a function of Gd-EOB-DTPA treatment time was fit into a logistic growth curve (R^2^=0.98). Its slope reached a maximum at 32.9 min, after which, uptake began to decrease (**Figure 2d**). No significant difference was observed between *R*_1_ at the 60-minute (5.03 ± 0.28 Hz) and 90-minute timepoints (5.67 ± 0.32 Hz; n=3, p=0.053) as the first derivative approaches zero, suggesting that the intracellular Gd(III) concentration detected via magnetic resonance approaches a steady state (*R*_1,max_= 5.85 Hz) at about 90 min of incubation (96.9 ± 5.5% of *R*_1,max_; **Figure 2d**). The slow uptake kinetics observed here as well as the slow cellular efflux of the probe (**Supplementary Figure S4**) reinforces the approach of previous work utilizing *oatp1a1*, wherein signal-to-background in mice reached a maximum at approximately 5 h post Gd-EOB-DTPA administration^23,25^.

Next, Luc-CTL and Luc-1B3 cells treated with Gd-EOB-DTPA were combined at various cell number ratios and their spin-lattice relaxation rates were measured at 3T. Relative to a pure sample of Luc-CTL cells (*i.e*. 0% Luc-1B3; 0.365 ± 0.013 Hz), the minimal volume fraction of Luc-1B3 cells required for a significant increase in *R*_1_ relaxation rate was 15% (0.96 ± 0.26 Hz; n=3, p=0.017; **Figure 2e**). A strong positive linear correlation between *R*_1_ relaxation rate and Luc-1B3 percent fraction was observed (n=3, R^2^=0.98; **Figure 2e**). In combination with the *R*_1_ signal-to-background ratios measured above, the results of these fractional 1B3 volume studies supported the feasibility of detecting engineered cell populations in animals with sensitivity at high resolution (*e.g*. 100-μm isotropic resolutions).

### *Oatp1b3* does not affect overall metastatic burden in mice and significantly enhances visualization of primary tumors and metastatic lesions *in vivo*

Using the results from the *in vitro* characterization experiments, we optimized our MRI parameters for *in vivo* longitudinal imaging of a spontaneous metastasis model of breast cancer in mice (**Figure 3**). In this model, as the primary tumor grows, metastatic lesions were expected to form across multiple lymph nodes and the lungs, in accordance with endpoint histology from previous literature (**Figure 3a**)^38^. It is worth noting that the temporal pattern of spread was not yet established prior to our study, but that the chronology reflected in **Figure 3a** is evidenced by longitudinal *oatp1b3*-MRI data presented in later figures. Nod scid gamma (NSG) mice were implanted with 3 × 10^5^ Luc-CTL (N=7) or Luc-1B3 (N=11) MDA-MB-231 cells at the left 4^th^ mammary fat pad and imaged via BLI (150 mg/kg D-luciferin) every two days, and pre- and post-contrast *T*_1_-weighted MRI at 3 T (1.3 mmol/kg Gd-EOB-DTPA) were acquired on day 12 for the primary tumor and day 26 for the upper body. Complete details on MRI parameters can be found in the *Materials and Methods*.

**Figure 3.**
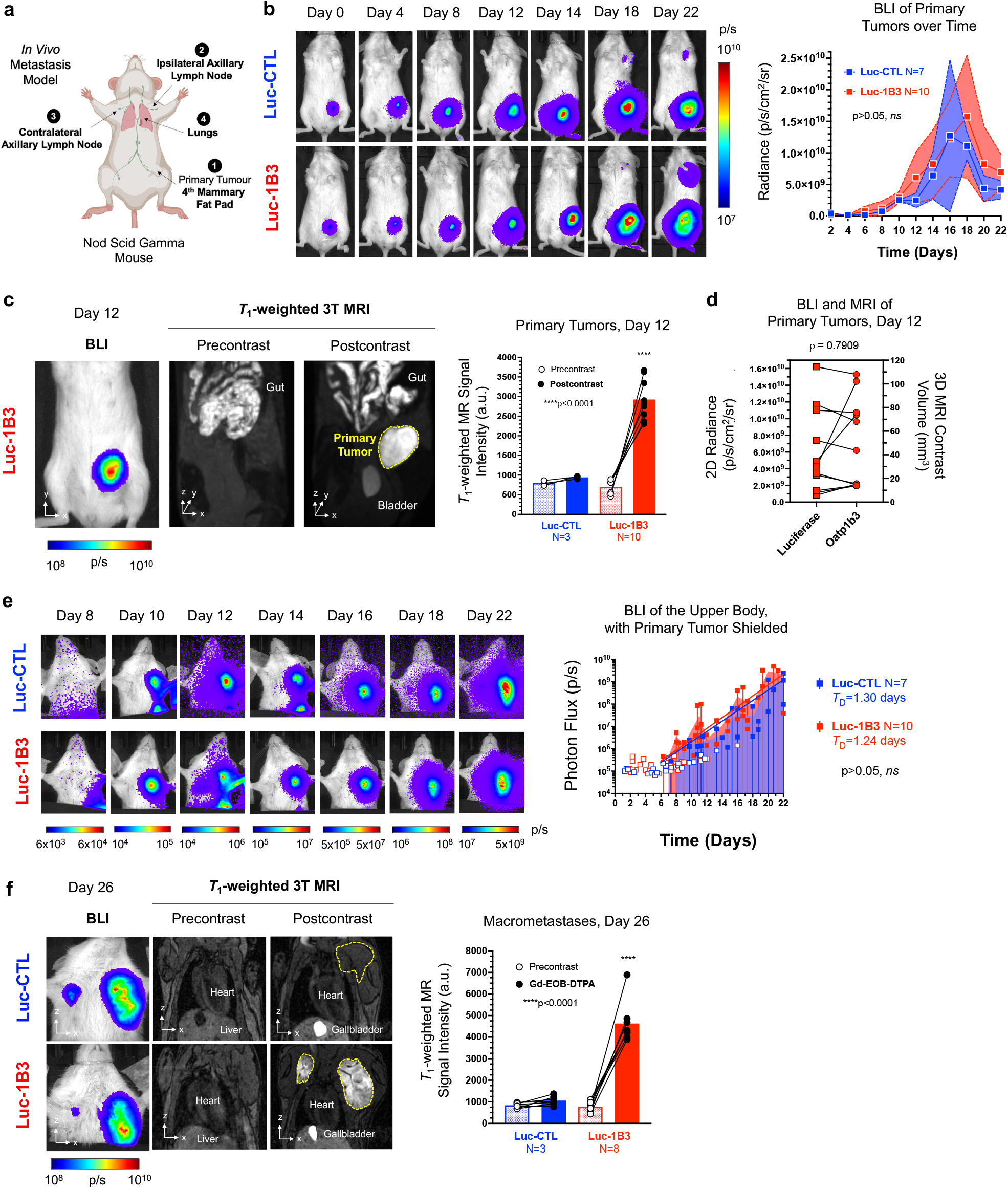
Deep Tissue Imaging of Primary Tumors and Metastatic Lesions via *Oatp1b3-MRI*. a) Spontaneous metastasis model. MDA-MB-231 cells are implanted orthotopically into the left-bearing 4^th^ mammary fat pad of a nod scid gamma mouse. Over time, the cells metastasize to the ipsilateral axillary lymph node, to the contralateral axillary lymph node, and to the lungs. b) Radiance (p/s/cm^2^/sr) from Luc-CTL (N=7, *blue*) and Luc-1B3 primary tumors (N=10, *red*) over time. p>0.05, ns. c) Same-day bioluminescence images (BLI), pre- and post-contrast *T*_1_-weighted MR images of representative Luc-1B3 mouse 12 days post cell implantation at the primary tumor site. Contrast enhancement of primary tumor is outlined in yellow. Average *T*_1_-weighted signal intensity (a.u.) of Luc-CTL (N=3, *blue*) and Luc-1B3 (N=10, *red*) primary tumors before and after Gd-EOB-DTPA administration, ****p<0.0001. d) 2D radiance (p/s/cm^2^/sr) and corresponding contrast-enhanced 3D volume (mm^3^) of individual Luc-1B3 tumors 12 days post cell implantation, n=10. Spearman rank coefficient, ρ=0.79, ***p=0.0085. e) Photon flux (p/s) measurements from the upper body of Luc-CTL (N=7, *blue*) and Luc-1B3 (N=10, *red*) mice over time. Hollow symbols represent signals from BLI with no discernible metastatic foci, whereas filled symbols represent photon flux measurements during times where at least one lesion was detected. *T*_D_, doubling time. p>0.05, ns. f) BLI, pre- and post-contrast *T*_1_-weighted images of the upper body of representative Luc-CTL and Luc-1B3 mice 26 days post cell implantation at the primary tumor site. Average *T*_1_-weighted signal intensity (a.u.) of Luc-CTL (N=3, *blue*) and Luc-1B3 (N=8, *red*) macro-metastatic lesions before and after Gd-EOB-DTPA administration, ****p<0.0001.

We first wanted to assess whether the *oatp1b3* reporter gene system interfered with primary tumor growth or metastatic progression. BLI demonstrated that primary tumors grew in both Luc-CTL (N=7) and Luc-1B3 (N=10) burdened mice, with no significant difference (p>0.42) in bioluminescent average radiance (p/s/cm^2^/sr) at each timepoint between the groups (**Figure 3b**). As primary tumors grew, BLI for metastatic lesions was performed by blocking light from the lower body to enable detection of smaller populations of engineered cells that may have metastasized to the upper body. No significant difference was observed in upper body photon flux (p/s) over time between Luc-CTL (N=7) and Luc-1B3 (N=10) mouse groups (p=0.18; **Figure 3e**), suggesting that *oatp1b3* reporter gene expression neither inhibited nor promoted metastasis. The doubling time of metastasis in the upper body for Luc-CTL mice was determined to be 1.30 ± 0.53 days compared to 1.24 ± 0.59 days for Luc-1B3 mice (p=0.83; **Figure 3e**).

On day 12, pre-contrast and post-contrast *T*_1_-weighted MRI at 3 Tesla exhibited an average 4.2-fold significant increase in signal intensity in Luc-1B3 primary tumors (p<0.0001; **Figure 3c**), whereas Luc-CTL primary tumors showed no difference between pre- and post-contrast images (1.2-fold, p>0.96; **Supplementary Figure S5**). There was a strong positive correlation between 2D radiance measurements from BLI and 3D contrast enhancement volumes generated via *oatp1b3*-MRI, with a Spearman rank-order correlation coefficient (ρ) of 0.791 (n=9, p=0.0085; **Figure 3d**) and a linear regression correlation coefficient (R^2^) of 0.56 (n=9, p=0.0055; **Supplementary Figure S6**). However, it is worth noting that the two systems were not absolutely correlated on day 12, even when necrosis was not yet a factor, as indicated by the homogeneity of contrast enhancement across all tumors on MRI (**Figure 3d**). On day 26, large contrast-enhanced metastatic lesions in the upper body were observed in Luc-1B3 mice post contrast (N=8), but not in Luc-CTL mice (N=3) following administration of Gd-EOB-DTPA (**Figure 3f**). Interestingly, the post-contrast signal intensity of the metastatic masses on day 26 (4630 ± 970 a.u.) was significantly greater than that of the pre-necrotic primary tumor volume on day 12 (2930 ± 510 a.u.; p<0.0001; **Figure 3f**).

### Imaging of spontaneous metastasis in single animals over time via *oatp1b3*-MRI reveals that BLI largely obscures detection of small, late-stage, and/or deep-seated lesions in mice

With the finding that the *oatp1b3* reporter gene system did not disable or promote metastatic progression, as evidenced on BLI, and the auxiliary finding of greater contrast enhancement from metastatic masses compared to their concomitant primary tumors, we then explored the system’s capability of longitudinally imaging the metastatic cascade via *oatp1b3*-MRI with a second cohort of mice (N=7; **Figure 4**). We wanted to determine the first timepoint at which metastatic lesions were detectable with either BLI or MRI. Mice were imaged according to the algorithm outlined in **Supplementary Figure S7**. Most mice were imaged up to day 22, but smaller subsets were imaged at timepoints beyond day 22, including day 26 (N=3), day 30 (N=3) and day 36 (N=3), the endpoint of which was determined by the health of each individual mouse. For five out of the 7 mice, metastatic lesions in the ipsilateral axillary lymph node were detected on MRI at the same timepoint as BLI (**Figure 4a**) whereas two mice exhibited signals on MRI at this site up to 48 h prior to BLI signal being detected (**Supplementary Figure S8**). Significantly increased MR signal intensity (2900 ± 620 a.u.) was exhibited relative to surrounding muscle tissue (1120 ± 110 a.u.; SBR = 2.59) at the time of detection (n=7, p<0.0001) (**Figure 4b**). On average, spontaneous metastases were first detected 11.2 ± 1.3 days post primary tumor implantation via *luciferase-BLI*, and 10.2 ± 0.8 days via *oatp1b3*-MRI (n=7, p>0.05, ns).

**Figure 4.**
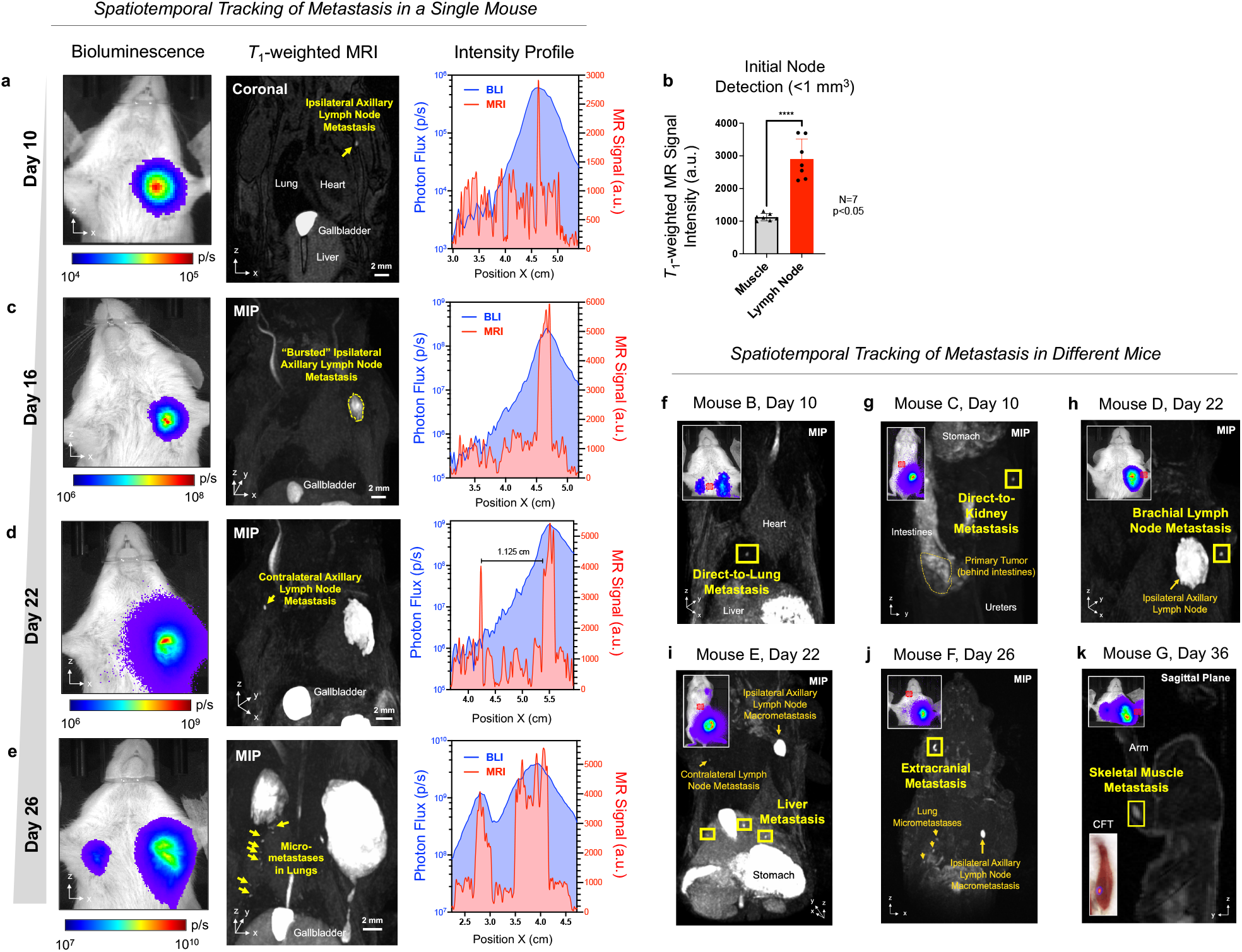
High Resolution, 3-Dimensional Tracking of Spontaneous Metastasis over Time. Representative bioluminescence images (BLI, p/s), post-contrast *T*_1_-weighted magnetic resonance images (MRI), and line profiles of BLI photon flux (p/s, *blue*) and MRI signal intensity (a.u., *red*) at 3T of one female nod scid gamma mice (n=7) after implantation of MDA-MB-231 cells into the left-bearing 4^th^ mammary fat pad on (a) Day 10, (c) Day 16, (d) Day 22, and (e) Day 26. MIP, maximal intensity projection. Scale bar, 2 mm. b) *T*_1_-weighted signal intensity (a.u.) at initial detection of ipsilateral axillary lymph node lesions (Days 10-12) and mean signal intensity (a.u.) of surrounding muscle tissue at 3 Tesla (n=7, ****p<0.0001). (f-k) Representative BLI and post-contrast *T*_1_-weighted MR images of unique patterns of metastasis across different mice on different days. Red square on BLI images indicates location of lesion determined by corresponding *oatp1b3*-MRI data. MIP, maximal intensity projection. CFT, cryo-fluorescence tomography.

The most prominent spatiotemporal pattern of metastasis that followed was continued growth of the initial metastatic lesion at the ipsilateral axillary lymph node, which was observed with both *luciferase-BLI* and *oatp1b3*-MRI (**Figure 4c**). Then, on day 22, all mice exhibited a significant increase in signal intensity on *T*_1_-weighted images at the contralateral axillary lymph node (2650 ± 790 a.u.; SBR = 2.44). These contralateral lesions could not be resolved on BLI until day 25.4 ± 1.3 due to light scatter from the first metastatic lesion on the opposite side of the animal (Δ*x*_avg_ = 1.14 ± 0.08 cm; **Figure 4d**). Following this, *oatp1b3*-MRI revealed that 6 of the 7 animals developed numerous micro-metastatic lesions (<1 mm^3^) in the lungs (**Figure 4e**). Again, these lung lesions were small in size and proximal to the larger lymph node lesions, thereby preventing their detection with BLI (**Figure 4e**).

Beyond this pathway, however, there was marked divergence in spatiotemporal patterns of metastasis between mice, even at early timepoints. Direct-to-lung metastasis was observed in one mouse on day 10, as both the signal intensity and volume of the lesion in the lung (*SI* = 3460 ± 260 a.u., *V* = 0.067 mm^3^) was already greater than that of its ipsilateral axillary lymph node lesion (*SI* = 2510 ± 120 a.u., *V* = 0.015 mm^3^; **Figure 4f**); because of the lesion’s location behind the heart, as indicated by *T*_1_-weighted images, BLI signals presented as diffuse and relatively imprecise (**Figure 4f**). In another case, a metastatic lesion in the contralateral kidney was detected via *oatp1b3-MRI* on day 10 (*SI* = 5670 ± 320 a.u., *V* = 0.071 mm^3^, d = 1.39 cm, SBR = 2.84; **Figure 4g**), but was not detected on BLI due to confounding signals stemming from the primary tumor (Δ*x* = 1.66 cm). Two additional mice exhibited metastasis to the kidneys on MRI at later dates, which again, could not be resolved on BLI.

On day 22, a minority of mice (3/7) exhibited continued spread through the lymphatic system to the ipsilateral brachial node on *T*_1_-weighted images, but this was virtually impossible to detect on BLI due to the close proximity of the node (5.9 mm in one representative mouse) to the initial metastatic lesion (**Figure 4h**). In one mouse, liver metastases (*V*_total_ = 0.18 mm^3^) appeared and outpaced its lung metastases (*V*_total_ = 0.025 mm^3^; **Figure 4i**). In another case, an extracranial metastasis in the head was detected via *oatp1b3*-MRI, but was obscured on BLI, likely due to the depth of the lesion (d = 5.6 mm, *V* = 0.15 mm^3^), relative to the larger, more superficial lymph node metastasis (d = 1.8 mm, *V* = 0.68 mm^3^), in combination with its cavitation between bones in the head, as indicated by *T*_1_-weighted images (**Figure 4j**). Finally, one mouse also exhibited metastasis to skeletal muscle (4920 ± 970 a.u., 0.43 mm^3^, SBR = 3.9) within its ipsilateral front leg, which was again, undetected on BLI due to its proximity to the initial lesion at the ipsilateral axillary lymph node (**Figure 4k**).

### *Oatp1b3*-MRI enables detection of metastatic lesions comprising fewer than 10^3^ cells on high resolution, 3-dimensional images of live mice

An analysis of mean signal intensity and volume of individual metastatic foci demonstrated that, for 3 mice, 60 micro-metastatic lesions with significantly higher MRI signal than lung background were detected in the lungs at about 30 days post cell implantation (**Figure 5b, 5c**). None of these lesions were detectable with BLI. The smallest volume that demonstrated mean signal intensity (3330 a.u.) on *T*_1_-weighted images considerably above the mean intensity of lungs in control animals (240 ± 170 a.u.), was measured at 0.0043 mm^3^ with image rendering (5 voxels without rendering), but a more reproducible minimal volume of detection was highlighted by a cluster of lesions (N=7) at around 0.01 mm^3^ (9-12 voxels; **Figure 5b**). For an epithelial tumor cell line, literature values equate a volume of 1 cm^3^ to approximately 10^8^ cells^39^. Back-calculating from this conversion, 0.01 mm^3^ would equate to approximately 10^3^ cells per lesion, or 10^2^ cells per voxel, assuming that these voxels were comprised entirely of Luc-1B3 cells. However, it is also worth noting that the range of signal intensities across these foci (1778 ≤ *SI* ≤ 4813 a.u.) did not correlate with lesion size (*R*^2^=0.059, p=0.13), indicating that many of these micro-metastatic volumes were not comprised entirely of Luc-1B3 cells, and the actual detection limit for *oatp1b3*-MRI likely falls below 10^3^ cells per lesion.

**Figure 5.**
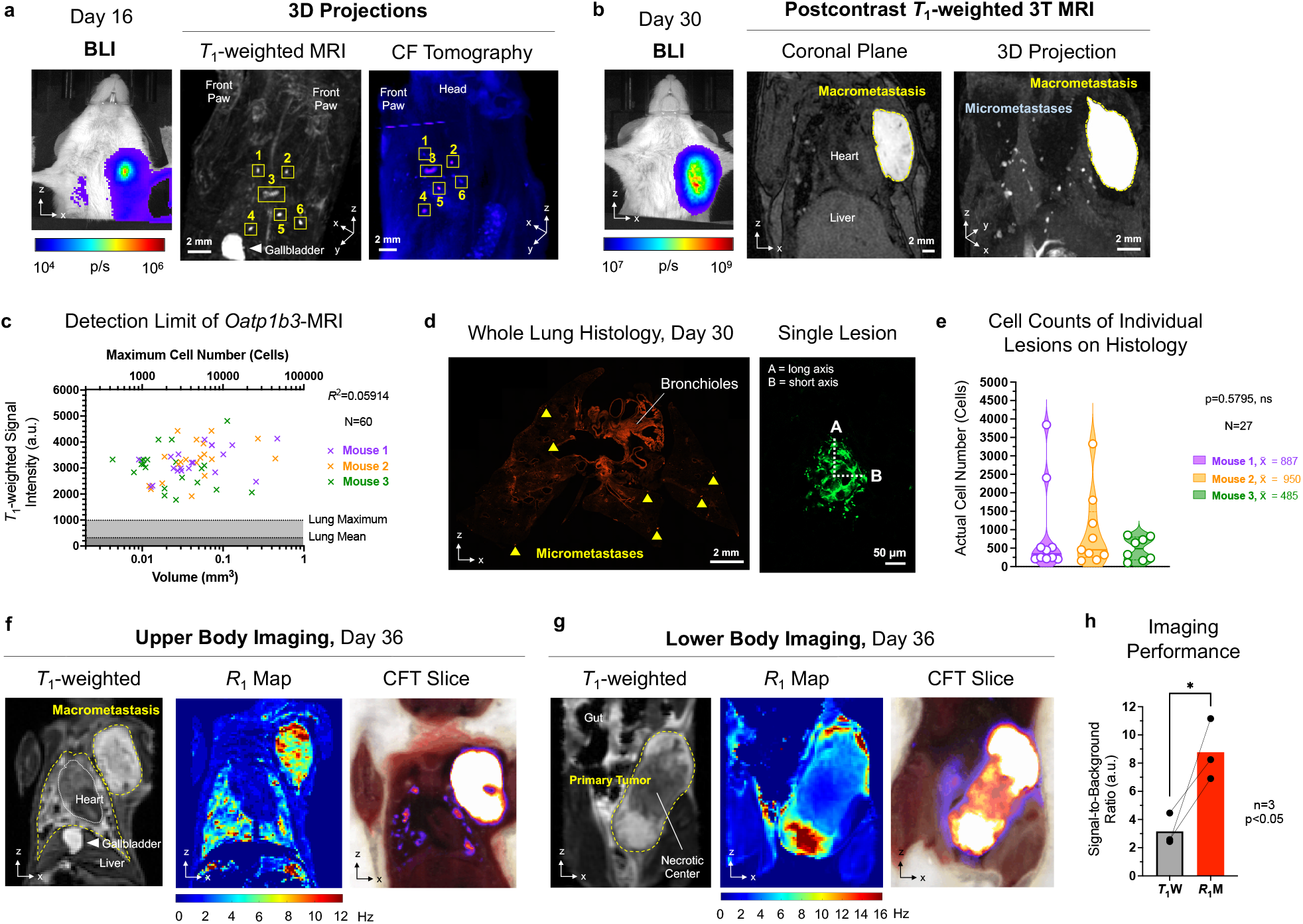
Ultrasensitive Detection of Micrometastases in Live Animals. a) Bioluminescence image (BLI, p/s), raw 3D projection of post-contrast *T*_1_-weighted MRI, and raw 3D projection of cryo-fluorescence tomography (CFT) of a Luc-1B3 mouse burdened with lung micrometastases that was imaged and sacrificed 16 days post implantation of its primary tumor. Numbered squares on projections indicated matched MR to CFT lesions. Scale bar, 2 mm. b) BLI (p/s), post-contrast *T*_1_-weighted 3T image, and maximum intensity projection of representative Luc-1B3 mouse on day 30 burdened with a unilateral macro-metastatic lesion and numerous micro-metastatic lesions across the lungs. Scale bar, 2 mm. c) *T*_1_-weighted signal intensity (a.u.) and volume (mm^3^) of individual micro-metastatic foci (N=60) across Luc-1B3 mice on day 30. n=3, *R*^2^=0.059. d) Representative whole-lung histological section of Luc-1B3 mouse shown in (a), imaged via red fluorescence for detection of engineered cells. Metastatic cell populations are indicated by yellow arrows. Scale bar, 2 mm. Note that bronchioles exhibited autofluorescence. Representative high magnification image of single metastatic lesion. Scale bar, 50 μm. e) Cell counts from individual lesions on histology across Luc-1B3 mice (N=27). Mean cell number per lesion, 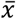. p>0.05, ns. Representative *R*_1_-weighted image (a.u.), *T*_1_ relaxation rate map (Hz) and cryo-fluorescence tomography (CFT) section (a.u.) of Luc-1B3 mice on Day 36. One mouse exhibits a large macro-metastasis and significant lung metastasis (f), and a second exhibits a primary tumor with a large necrotic core (g). h) Average signal-to-background ratio (SBR, a.u.) of post-contrast Luc-1B3 lesions on *T*_1_-weighted images (*T*_1_W), and *R*_1_ maps (*R*_1_M), n=3, p<0.05.

We attempted (N=1) to match micrometastases in the lungs detected on MRI to corresponding lesions on whole-mouse histology generated via the Xerra™ imaging system (EMIT Imaging, Baltimore, MD). Coordinates were conserved between the two 3D datasets in some localized regions (**Figure 5a**), but the lack of structural reinforcement in the lungs led to considerable architectural deformation during histological processing, such that matching lesions from MR images to their precise locations on histology proved challenging without non-rigid image registration that would disturb image integrity considerably. Instead, to acquire a proximal ground truth measurement of detection sensitivity, mice imaged on day 30 were immediately sacrificed and the number of cells per micro-metastatic lesion in lung tissue was extrapolated from section microscopy (**Figure 5b–5e**). These measurements supported our *in vivo* calculations, as well as the hypothesis that the *oatp1b3*-MRI system can detect populations comprising fewer than 10^3^ cells in lung tissue. Five lesions across three mice comprised of greater than 10^3^ cells, with the largest measured at an estimated 3840 cells, but all remaining lesions fell below this threshold. Overall, on average, micro-metastases in lung tissue on day 30 were estimated to be comprised of 789 ± 185 cells (n=3 mice, N=27 lesions). As assurance, and in agreement with our MR images, no macro-metastatic lesions (>1 mm^3^) were detected in lung tissue histology of these mice.

Finally, we sought to employ quantitative MRI methods to possibly improve *oatp1b3*-MRI performance even further. We reasoned that measuring *R*_1_ relaxivity (Hz) *in vivo* would generate greater signal-to-background ratios for more sensitive and specific detection of engineered cells, relative to signal intensity (a.u.) measurements generated from *T*_1_-weighted image data. To obtain 3D *R*_1_ maps of mice in a practical timeframe (10-15 minutes per mouse), we established a 4-flip-angle pulse sequence pipeline and postprocessing algorithm based on the driven equilibrium single pulse observation of *T*_1_ (DESPOT1) method^40^. During imaging, *T*_1_-weighted images were also acquired at comparable acquisition times and equivalent resolutions for each mouse subject (n=3, **Figure 5f, 5g**). Complete details on MRI parameters are outlined in the *Materials and Methods* section. Although previously recorded values for *R*_1_ rates of non-brain murine tissues at 3T are relatively sparse, our *in vivo* measurements for healthy control tissues, *e.g*. heart, skeletal muscle, fat, were in general agreement with previously published data^41^. Importantly, the SBR of Luc-1B3 lesions on *R*_1_ maps was significantly greater (8.8 ± 2.2) than the SBR of the same lesions on *T*_1_-weighted images (3.3 ± 1.1, p=0.017, **Figure 5h**).

## DISCUSSION

Tracking spatiotemporal patterns of metastatic spread in deep tissues of single animals is a critical experimental capability for cancer research. Yet, sensitive and quantitative preclinical assessment of metastatic disease with a high degree of accuracy remains a significant challenge. Compared with traditional contrast agents, reporter genes offer information on cell viability, and do not require the presence of biomarkers to be specific to the cells of interest, as is needed to study triple-negative breast cancer and many other cancer subtypes. To overcome the intensiveness and sampling bias of traditional histology, the last decade has seen a surge of tissue clearing methods that render large biological samples transparent and allow three-dimensional views of large tissue volumes^42^. Still, a major drawback of this approach, as with histology, is that only a single timepoint can be acquired of an individual animal, which in turn, necessitates processing of large numbers of animals to draw meaningful conclusions^43^. This is an especially notable issue when working with spontaneous metastasis models that exhibit high variability between animals with respect to both the rate and location of metastasis^44^.

BLI avoids single-timepoint limitations by providing sensitive whole-body information on relative locations of cancer cells in live mice, but we demonstrate that smaller cell populations largely go undetected due to the presence of larger lesions. Specifically, we show that small, single lesions positioned at distances >1 cm from larger engineered cell populations go undetected unless they grow to substantial sizes before endpoint, and that populations of micro-metastases in separate organs remain obscured throughout the BLI timeline (**Figure 4**). In summary, BLI data is skewed towards initial, superficial lesions, that do not progress and respond to treatment in the same way as smaller, late-stage, and/or deep-seated metastases located in distinct micro-environments^45^. In fact, promotion of metastasis by antitumor therapies has been previously demonstrated in several major studies^46–48^. And on the opposite end, it may be that successful treatments for metastasis have been wrongly deemed ineffective simply because larger lesions dominated measurements^49^. Yet, above all, it is concerning that little emphasis has been placed on the discovery of antimetastatic therapies at the preclinical stage because the threshold of metastatic prevention and/or regression is extremely challenging to demonstrate with currently available technologies^50,51^.

*Oatp1b3*-MRI overcomes many limitations for accurate spatiotemporal tracking of metastatic spread *in vivo*. We show that this system can dynamically track the metastatic process in its earliest stages, at small lymph node lesions, even prior to BLI detection in some cases. At late stages, we demonstrate its superiority over BLI to track cancer spread to multiple lymph nodes and other deep-seated organs on highly-resolved images, owing to the inherent absence of signal scatter and signal attenuation in MRI (**Figure 4**). In parallel, *oatp1b3*-MRI enabled detection of metastatic lesions with high sensitivity, exhibiting an *in vivo* threshold on the order of less than 10^3^ cells per lesion, or 10^2^ cells per voxel, in the lungs (**Figure 5**), signifying a 12.5-fold improvement in minimum detection volume relative to the latest state-of-the-art MR methods^15^. Although the sites of metastasis in this study are in close agreement with previously published literature^38^, and a pattern of metastasis similar to that observed in human breast cancer patients is exhibited^52^, detailed tracking of metastatic spread in single animals from one location to the next has not previously been demonstrated. Critically, imaging mice at multiple timepoints with *oatp1b3*-MRI did not result in differences in metastatic spread relative to control mice, making it suitable for future studies focused on assessing antimetastatic therapies *in vivo*.

However, the *oatp1b3*-MRI system is itself not without limitations, and room still remains to improve its imaging capabilities with respect to both specificity and sensitivity. On the matter of whole-body imaging, the stomach and intestines exhibited high signal intensities on both precontrast and postcontrast *T*_1_-weighted images, which may have obscured detection of small metastases at these sites; in addition, the small molecule probe Gd-EOB-DTPA does not readily cross the blood-brain barrier (BBB), hindering detection of brain metastasis until lesions grow enough to compromise BBB permeability. Although these issues can be addressed through setups that enable pre/postcontrast-image subtraction and focused-ultrasound BBB opening, respectively, efforts are warranted to fully streamline *oatp1b3*-MRI. Simply but effectively, incorporating a potato diet for 24h before imaging has been shown to practically eradicate non-specific gastrointestinal *T*_1_-weighted signals in mice and should therefore be implemented into all future *oatp1b3*-MRI animal protocols^53^.

Additionally, new paramagnetic probes targeting *oatp1b3* are underway, and small lipophilic molecules would be prime candidates to unlock the capability of sensitively detecting brain metastasis on *oatp1b3*-MRI with the BBB still intact^54,55^. However, one outstanding blind spot with little room for mitigation is the gallbladder, as it represents a collection site for *oatp1b3*-targeted probes that are categorically eliminated via the hepatobiliary pathway; but, we believe this represents a minor drawback in the context of the larger system. Finally, protein engineering provides opportunities to further improve system parameters and functionalize *oatp1b3* as a biosensor. For example, removal of the protein’s downregulatory phosphorylation sites may increase the steady state concentration of OATP1B3 at the cell membrane for greater probe influx capacity; directed evolution can be employed to screen for mutants with rapid probe-transport kinetics; and, addition of sensory motifs can be added to enable imaging of events like deep-brain neuronal activation. For now, however, *oatp1b3*-MRI offers a means to track *oatp1b3*-engineered cells in deep tissues of live animals over time with combined high resolution and high sensitivity. We anticipate that this platform will facilitate our abilities to cultivate clinically-predictive preclinical models of metastasis^56^, expand our understanding of the metastatic process, and provide a means to rigorously evaluate antimetastatic therapies *in vivo*.

## MATERIALS AND METHODS

### Lentivirus Construction and Production

Lentiviral transfer plasmids co-encoding tdTomato fluorescent protein with human codon-optimized firefly luciferase 2^24^, and zsGreen fluorescent protein with Oatp1b3^57^ were cloned (**Figure 1b**). The cDNA for Organic anion transporting polypeptide 1b1 (Oatp1b1) was acquired from the hOATP1B1/SLCO1B1 VersaClone cDNA Vector (Cat. RDC1113, R&D Systems, Minneapolis, Minnesota, United States). A lentiviral transfer plasmid co-encoding zsGreen with Oatp1b1 was cloned using the In-Fusion HD Cloning kit (Takara Bio USA Inc, Madison, Wisconsin, United States). Third-generation packaging and envelope-expression plasmids (pMDLg/pRRE, pRSV-Rev, and pMD2.G, Addgene plasmids: #12251, #12253, and #12259, respectively; gifts from Didier Trono) were co-transfected with each of the three transfer plasmids (tdTomato/Luciferase, zsGreen/Oatp1b1, zsGreen/Oatp1b3) into human embryonic kidney (HEK 293T) cells using Lipofectamine 3000 according to the manufacturer’s lentiviral production protocol (Thermo Fisher Scientific Inc., Waltham, Massachusetts, United States). Lentivirus-containing supernatants were harvested 24h and 48h post-transfection, filtered through a 0.45-μm filter, and used immediately for transductions.

### Cell Culture and Stable Cell Generation

HEK 293T and human triple negative breast cancer cells (MDA-MB-231) were obtained from a commercial supplier (American Type Culture Collection, Manassas, Virginia, United States) and cultured in Dulbecco’s Modified Eagle’s Medium (DMEM) supplemented with 10% fetal bovine serum at 37°C and 5% CO_2_. All cells were routinely verified as free of mycoplasma contamination using the MycoAlert mycoplasma detection kit (Lonza Group, Basel, Switzerland). MDA-MB-231 cells were first transduced with lentivirus (MOI=5) co-encoding tdTomato and human codon-optimized firefly luciferase 2, and fluorescence-activated cell sorting was performed to sort for tdTomato-positive cells using a FACSAria III cell sorter (BD Biosciences, Mississauga, Ontario, Canada). The sorted cells were expanded and transduced again with lentivirus (MOI=5) co-encoding zsGreen and Oatp1b3 and fluorescence-activated cell sorting was performed to select for tdTomato and zsGreen double-positive cells using equivalent intensity-gating for each colour. Sorted cells, referred to as either Luc-CTL or Luc-1B3 cells, were utilized for all subsequent experiments.

### *In Vitro* Bioluminescence

Luc-CTL and Luc-1B3 cells were seeded into 24-well plates with the following numbers of cells per well: 1×10^6^, 5×10^5^, 3×10^5^, 1×10^5^, and 5×10^4^. Immediately after seeding, 0.15 mg/ml D-luciferin was added to each well, and plates were imaged on an IVIS Lumina XRMS *In Vivo* Imaging System (PerkinElmer, Waltham, Massachusetts, United States). Average radiance values in p/s/cm^2^/sr were measured from each well using Living Image software (PerkinElmer, Waltham, Massachusetts, United States).

### Western Blot

1×10^6^ Luc-CTL or Luc-1B3 cells were washed three times in PBS and incubated with 200-μL of chilled RIPA buffer and protease inhibitors for 30 minutes. Lysates were collected and sonicated with five 5.0-s 40-kHz bursts before being centrifuged at 13,000g for 20 minutes at 4°C. Supernatants were collected, quantified and 40 μg of protein from each sample was loaded into an acrylamide gel composed of a 4.0% stacking layer buffered at pH 6.8 and a 15% separation layer buffered at pH 8.8. Gel electrophoresis was performed for 20 minutes at 90V and 1 hour at 110V. Protein was transferred to a nitrocellulose membrane for 7.5 minutes via the iBlot™ 2 Gel Transfer Device (IB21001, Thermo Fisher Scientific, Waltham, Massachusetts, United States) and blocked with 0.05% Tween-20, 3% BSA solution for 30 minutes. Rabbit anti-Oatp1b3 antibody (1:1000 dilution, ab139120, Abcam, Cambridge, United Kingdom) was added and incubated overnight at 4°C. The blot was washed 3× with 0.05% Tween-20 solution for 10 minutes and Goat anti-Rabbit 790-nm antibody (1:10,000 dilution, A11369, Thermo Fisher Scientific, Waltham, Massachusetts, United States) was added for 45 minutes at room temperature. The blot was washed again 3× with 0.05% Tween-20 solution for 10 minutes and imaged on the Odyssey CLx Imaging System (LI-COR Biosciences, Lincoln, Nebraska, United States).

### Transmission Electron Microscopy

3×10^5^ Luc-1B3 cells were seeded and grown on Thermanox cover slips (150067, Thermo Fisher Scientific, Waltham, Massachusetts, United States) until confluency was reached, after which, cells were incubated with 1 mM Gd-EOB-DTPA or an equivalent volume of saline in DMEM for 1 hour. Following 1 hour of incubation, cells were immediately washed 3× with PBS, and incubated for 10 minutes with a 1:1 solution of Karnovsky’s fixative (2% paraformaldehyde, 2.5% glutaraldehyde in 0.1 M cacodylate buffer, pH=7.4) and DMEM at 37°C, and then subsequently incubated for 3 hours with Karnovsky’s fixative at room temperature. Finally, cells were washed 4×, for 5 minutes each time, with 0.1M cacodylate buffer.

### Induced Coupled Plasma Mass Spectrometry

3×10^5^ Luc-CTL and Luc-1B3 cells (n=3) were seeded in 6-well plates and allowed to grow for approximately three days, until confluency was reached. Cells were then treated with variable concentrations of Gd-EOB-DTPA (0, 1.6, 3.2, 6.4, 9.6, 16.0 mM), washed 3× with PBS, trypsinized and counted. One million cells of each condition were harvested and lysed via 20-minute shaking incubation with 500 μL of 25 mM Tris•HCl, 150 mM NaCl, 1% NP-40, 1% sodium deoxycholate and 0.1% sodium dodecyl sulfate. The lysate was then digested by adding 10% (v/v) HNO_3_ solution to bring the final sample volume to 10 ml.

### *In Vitro* Nuclear Magnetic Relaxometry

For all *in vitro* magnetic characterization experiments, one million Luc-CTL or Luc-1B3 cells were seeded in T-175 cm flasks and allowed to grow for three days. Cells were then incubated with Gd-EOB-DTPA or Gd-DTPA at specific concentrations and lengths of time prior to being washed 3× with PBS and collected for imaging. The following parameters were used for each experimental objective:

- For NMRD, Luc-CTL and Luc-1B3 cells were incubated with 16 mM Gd-EOB-DTPA or Gd-DTPA for 1 hour (**Figure 2b**). For measurements at 3T, Luc-CTL and Luc-1B3 cells were incubated with either 1.6 or 16 mM Gd-EOB-DTPA or Gd-DTPA for 1 hour (**Figure 2c**).
- To evaluate reporter-probe kinetics, Luc-1B3 cells were incubated with 16 mM Gd-EOB-DTPA for 0, 10, 20, 30, 45, 60, or 90 minutes (**Figure 2d**). In parallel, Luc-CTL cells were incubated with 16 mM Gd-EOB-DTPA and Luc-1B3 cells were incubated with 16 mM Gd-DTPA for 90 minutes as experimental controls.
- To determine volume fraction sensitivity, Luc-CTL and Luc-1B3 cells were incubated separately with 16 mM Gd-EOB-DTPA for 1 hour, counted, and mixed into tubes at the following Luc-CTL to Luc-1B3 ratios: 1:0 (0%), 99:1 (1%), 195:5 (2.5%), 19:1 (5%), 9:1 (10%), 17:1 (15%), 4:1 (20%), 3:1 (25%), 1:1 (50%), and 0:1 (100%).

Following washing, cells were trypsinized, centrifuged and 10 million cells were counted and placed in 300-μL tubes. Nuclear magnetic relaxation dispersion data for the resulting cell pellets were acquired at magnetic fields from 230 μT to 1 T on a fast field-cycling NMR relaxometer (SpinMaster FFC2000 1T C/DC, Stelar, s.r.l., Mede, Italy) by changing the relaxation field in 30 steps, logarithmically distributed using an acquisition field of 380.5 mT, with temperature kept constant at 37°C. Data for each sample were subsequently fit into a LOWESS spline curve with 10 points set in the smoothing window.

*In vitro* MRI was performed on a 3-T GE clinical MR scanner (General Electric Healthcare Discovery MR750 3.0 T, Milwaukee, Wisconsin, United States) using a clinical 16-channel head birdcage RF coil. A mold was used to make a sample holder for the cell pellets comprised of 1% agarose that organized the tubes in a circular pattern to ensure equidistance of all samples to the coil during imaging and reduce magnetic susceptibility artefacts. Imaging data were acquired at 37°C with a fast spin-echo inversion-recovery (FSE-IR) pulse sequence using the following parameters: matrix size = 256 × 256, repetition time (TR) = 5000 ms, echo time (TE) = 19.1 ms, echo train length (ETL) = 4, number of excitations (NEX) = 1, receiver bandwidth (rBW) = 12.50 kHz, inversion times (TI) = 20, 35, 50, 100, 125, 150, 175, 200, 250, 350, 500, 750, 1000, 1500, 2000, 2500, 3000, in-plane resolution = 0.2 × 0.2 mm^2^, slice thickness = 2.0 mm, acquisition time = 5 min, 25 s per inversion time. Spin-lattice relaxation rates (*R*_1_) were determined by non-linear least-squares fitting (MATLAB, MathWorks, Natick, Massachusetts, United States) of signal intensities across the series of variable inversion time images on a pixel-by-pixel basis using the following standard model:

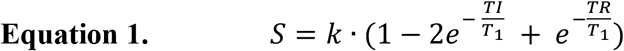

Here, *S* represents the acquired signal, and *k* is the proportionality constant, which depends on the specific coil used, the main magnetic field, the proton density, and the temperature, amongst other factors ^58^. However, two practical obstacles exist that render this equation impractical for real-world applications: first, a perfect 180° pulse is never truly achieved, and second, the signal is not acquired immediately after the RF pulse^59^. In practice, these real-world limitations produce a different behavior at *TI* = 0 and *TI* = ∞ than predicted theoretical outputs dictated by the equation above. To address these limitations, and assuming *TR* » *TI*, the equation above was adjusted to the following:

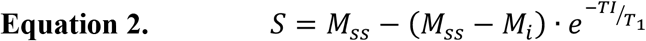

Here, if *TI* = 0, then *S* = −*M_i_* where *M_i_* is simply the first value of the inversion recovery curve, independent of the final steady state magnetization *M_ss_*. Additionally, if *TI* = ∞, then *S* = *M_ss_*. Finally, we use the absolute value of this last equation because the stored DICOM images acquired provided only magnitude (non-phase) information:

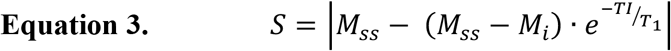

The zero-crossing point, the *TI* at which point *S* = 0, is extrapolated from the data and the *R*_1_ of each pixel can be determined via the following relationship:

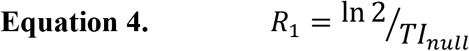

Source code is available upon request.

### Spontaneous Metastasis Model

Animals were cared for in accordance with the standards of the Canadian Council on Animal Care, and experiments were undertaken with an approved protocol of the University of Western Ontario’s Council on Animal Care (AUP 2016-026). Luc-CTL or Luc-1B3 cells (3×10^5^) were implanted into the left 4th mammary fat pad of female mice (NOD-*scid* IL2Rgamma^null^ strain, NSG, Jackson Laboratory, Bar Harbor, Maine, United States). BLI was performed after 150-mg/kg D-luciferin injection. *T*_1_-weighted 3T-MRI was performed before and 5h post 1.3-mmol/kg Gd-EOB-DTPA injection. Detectability of widespread metastases was evaluated by imaging Luc-CTL and Luc-1B3 mice (n=3 each) 30 days after cell implantation. Subsequently, a second Luc-1B3 cohort (n=7) was imaged over time until endpoint (Day 22-36, dependent on the health of each mouse) to assess ability to monitor metastasis dynamically.

### *In Vivo* Bioluminescence Imaging

BLI was performed on an IVIS Lumina XRMS *In Vivo* Imaging System (PerkinElmer). Mice were anesthetized with 1-2% isoflurane using a nose cone attached to an activated carbon charcoal filter for passive scavenging and administered 150 μL of 30-mg/mL D-luciferin intraperitoneally. Whole-body BLI was acquired with repeated 1.0-s exposure times every minute for approximately 15 minutes. Once the maximum signal plateaued, the lower half of the mouse *i.e*. location of primary tumor, was shielded with opaque black cloth and the front limbs of the mice were taped down to fully expose the anterior side of the thoracic region. The field-of-view was adjusted to fit the upper body of the mouse and an image was captured with a 5-minute exposure time to detect spontaneous metastases. Regions-of-interest (ROIs) were manually drawn around primary tumor borders using LivingImage software (PerkinElmer, Waltham, Massachusetts, United States) to measure bioluminescent average radiance (p/s/cm^2^/sr). For measurements of total metastatic burden in the upper body, rectangular ROIs were drawn according to the perimeter of the mouse thorax, and total photon flux (p/s/) from this region was calculated.

### *In Vivo* Magnetic Resonance Imaging

Mice (n=7) were anesthetized with 1–2% isoflurane by using a nose cone attached to an activated carbon charcoal filter for passive scavenging and positioned in a lab-built tray that was warmed to 40°C during MR imaging. All *in vivo* MR imaging used a clinical 3-Tesla GE MR750 clinical scanner (General Electric Healthcare, Milwaukee Wisconsin, USA) with a custom-built insert gradient: inner diameter=17.5 cm, gradient strength = 500 mT/m, peak slew rate = 3000 T/m/s; and a bespoke 3.5-cm diameter, 5.0-cm length birdcage radiofrequency coil (Morris Instruments, Ottawa, Ontario, Canada). Pre-contrast *T*_1_-weighted images were acquired using a three-dimensional Fast Spoiled Gradient Recalled Acquisition in Steady State (FSPGR) pulse sequence using the following parameters: FOV = 40 mm, TR = 12.0 ms, TE = 3.2 ms, rBW = 31.25 kHz, ETL = 4, Matrix size 400×400, Flip Angle = 60°, NEX = 3, voxel size = 100-μm isotropic, scan time = 18–24 minutes per mouse, dependent on mouse size. Mice were intravenously injected with 1.3 mmol/kg Gd-EOB-DTPA and imaged again approximately 5 hours post injection. Volumes-of-interest (VOIs) were manually delineated around metastatic lesions in post-contrast images using free and open-source code ITK-SNAP^60^ (www.itksnap.org) and Horos Project software (horosproject.org, Nimble Co LLC d/b/a Purview, Annapolis, Maryland, United States). VOIs were compared on pre-contrast and post-contrast images for differences in image contrast.

For quantitative *R*_1_ mapping, the driven equilibrium single pulse observation of *T*_1_ (DESPOT1) approach^40^ was employed. Successive images were acquired using the FSPGR sequence as described above at 5°, 10°, 15°, and 20° flip angles using an FSPGR pulse sequence: frequency FOV = 4.0 cm, phase FOV = 2.6 cm, slice thickness = 0.4 mm, TR = 4.5 ms, TE = 1.2 ms, matrix size = 100×100, number of excitations = 5, receiver bandwidth = 31.25 kHz, acquisition time = 2:27 to 2:50 minutes per flip angle (dependent on mouse size), total acquisition time = ~10-12 minutes per mouse. Pixelwise *in vivo* spin-lattice relaxation time maps for each timepoint were computed via MatLab (R2020a, MathWorks, Natick, Massachusetts, United States) by calculating the slope of the signal intensities of the raw images according to the following equation:

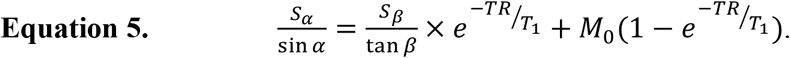

Here *S_α_* is the FSPGR signal intensity associated with flip angle *α*, *S_β_* is the FSPGR signal intensity associated with flip angle *β*, and *M*_0_ is the proportionality constant relating to the equilibrium longitudinal magnetization. Plotting 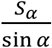 against 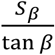 for each voxel allows for the determination of *T*_1_ from the slope *m* of this line as:

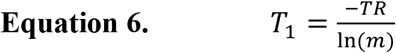

Source code is available upon request. All resultant images were reconstructed and analyzed on ITK-SNAP. Manual segmentation was performed for each mouse to create regions-of-interest for metastatic lesions as well as surrounding healthy tissue for calculations of signal-to-background ratios.

### Histology

Following *in vivo* experiments, mice were sacrificed via isoflurane overdose, perfused with 4% paraformaldehyde through the left heart ventricle, and the relevant organs were carefully excised. Tissues were then frozen in OCT medium (Sakura Finetek) and 10-μm or 150-μm frozen sections were collected onto glass slides. Whole-tissue microscopy images of tdTomato fluorescence were acquired using an EVOS FL Auto 2 Imaging System (Invitrogen) before hematoxylin/eosin (H&E) staining of the same and/or adjacent histology sections and subsequent imaging with the same microscope. For whole-mouse imaging, sacrificed mice were immediately submerged in hexanes over dry ice for flash freezing, and incubated at −80°C for ≥48 h prior to imaging on the Xerra™ system (EMIT Imaging, Baltimore, MD) at 60-μm isotropic resolutions.

### Statistical Analyses

Unless otherwise stated, statistical analysis was performed using Graphpad Prism software (Version 9.00 for Mac OS X; GraphPad Software Inc, La Jolla,CA; www.graphpad.com). Unpaired 2-tailed t-tests, and one or two-way analysis of variance (ANOVA) and Tukey post hoc multiple comparisons were performed, depending on the number of conditions and number of independent variables. For all tests, a nominal p-value less than 0.05 was considered statistically significant.

## ACKNOWLEDGEMENTS

Financial support for this manuscript was provided by Natural Sciences and Engineering Research Council of Canada (NSERC) Discovery Grants (JAR RGPIN-2016-05420, TJS RGPIN-2017-06338), and an Ontario Institute for Cancer Research Investigator Award (TJS IA-028). NNN is grateful to have received financial support from a Natural Sciences and Engineering Research Council of Canada Postgraduate Research Scholarship (2017-2021), and as an Amgen Awardee of the Life Sciences Research Foundation (2021-present). The authors would also like to acknowledge David Reese for providing helpful resources, and the larger Cellular and Molecular Imaging Group at the Robarts Research Institute for insightful discussions. The authors would also like to acknowledge Karen Nygard and Reza Khazaee of the Biotron Facilities at the University of Western Ontario for their expertise on transmission electron microscopy, as well as Patrick Zakrzewski, Mohammed Farhoud, and Mat Brevard from EMIT Imaging Technologies for their expertise in cryo-fluorescence tomography.

## CONFLICTS OF INTEREST

The authors declare no conflicts of interest.

---

**Figure S1.**
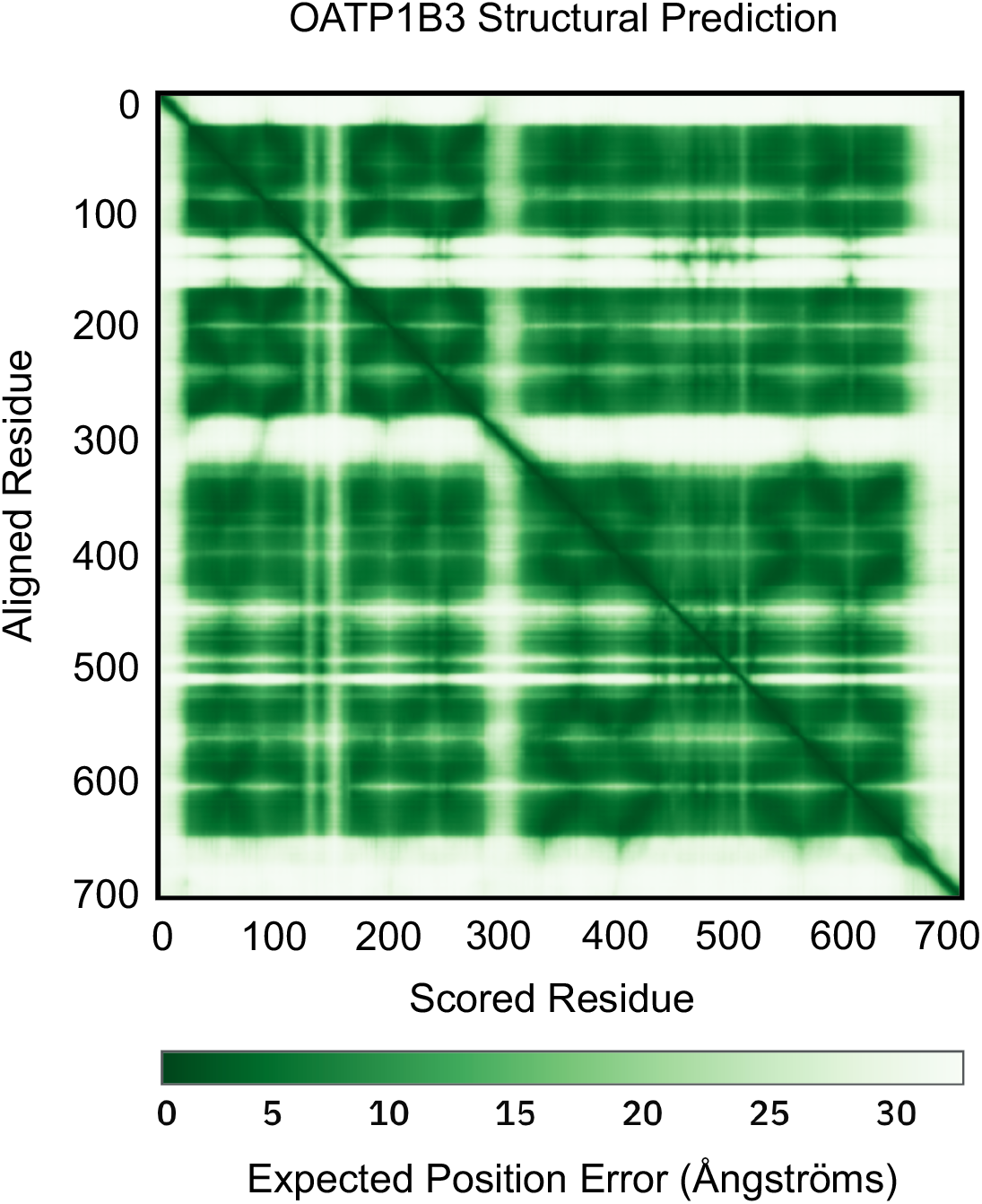
Predicted aligned error plot for OATP1B3, where expected position error for the aligned residue is assigned an expected position error (Ångströms) against the rest of the protein sequence. The transmembrane domains of OATP1B3 exhibit low expected position error (<5 Å) whereas high expected position error (>25 Å) is demonstrated at both termini, Extracellular Domain 2 (ECD2) and Cytoplasmic Domain 4 (CD4).

**Figure S2.**
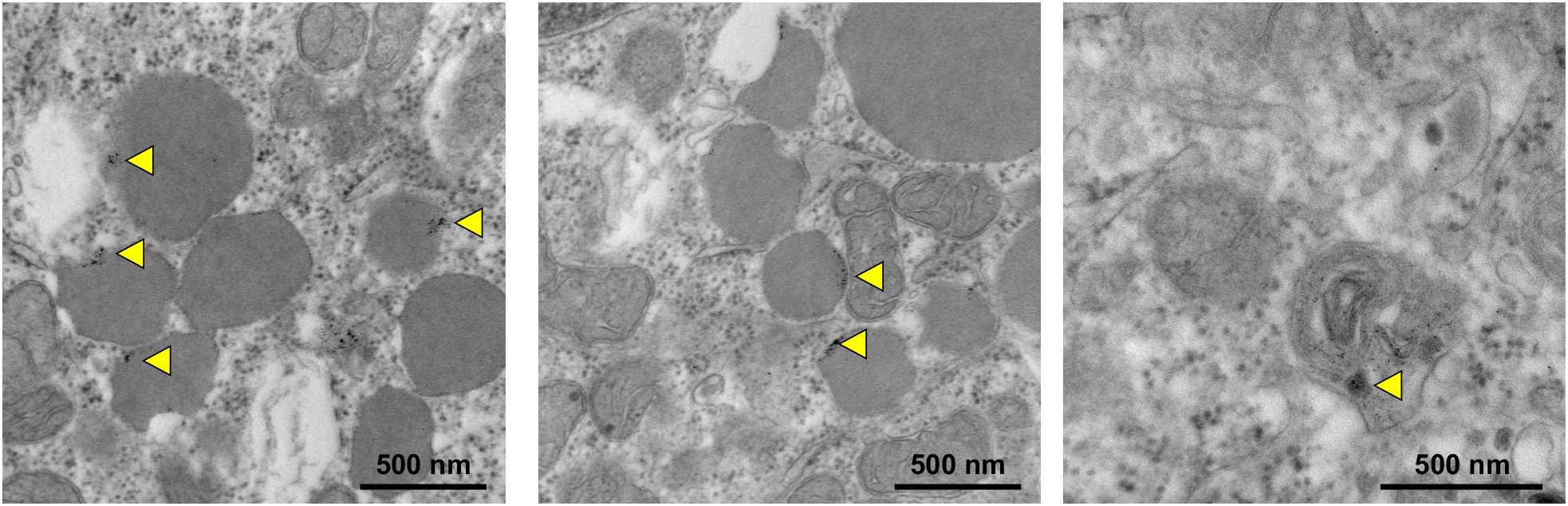
Transmission electron microscopy of Luc-1B3 cells incubated with 1 mM Gd-EOB-DTPA (appearing as black foci in TEM images, highlighted by yellow arrows) for 1 hour. Gd(III) is shown to be encapsulated in residual bodies, which are typically destined for exocytosis. Scale bar, 500 nm.

**Table S1.**
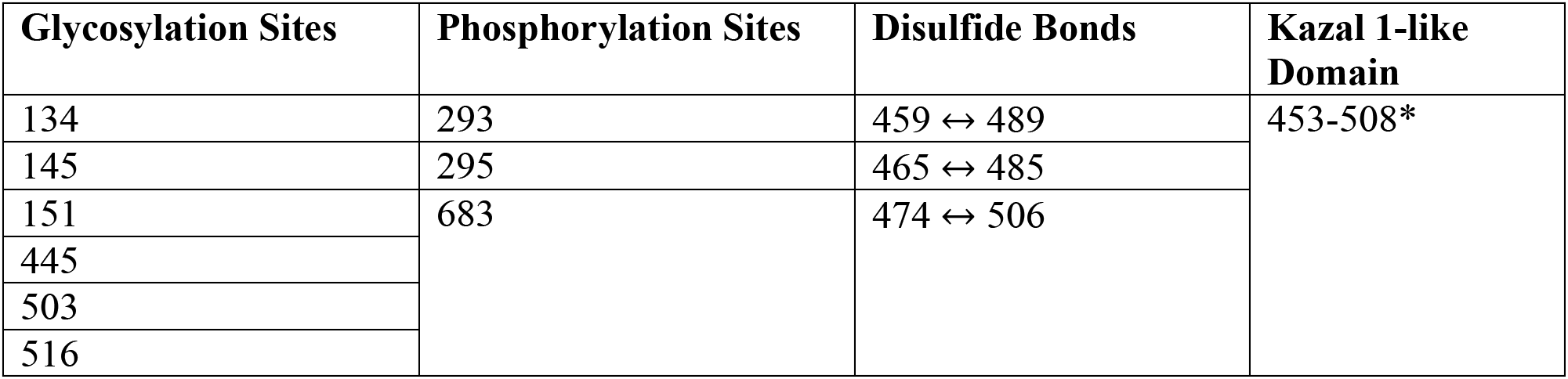
Predicted post-translational regulatory positions of OATP1B3. N-linked glycosylation to asparagine residues, which has been correlated to promotion of transporter function^31^. Phosphorylation of serine residues has been correlated to downregulation of transporter activity^29^. Disulfide bonds within the Kazal-1 like domain are predicted to result in serine protease inhibitor activity.

| Glycosylation Sites | Phosphorylation Sites | Disulfide Bonds | Kazal 1-like Domain |
| --- | --- | --- | --- |
| 134 | 293 | 459 ↔ 489 | 453-508* |
| 145 | 295 | 465 ↔ 485 |  |
| 151 | 683 | 474 ↔ 506 |  |
| 445 |  |  |  |
| 503 |  |  |  |
| 516 |  |  |  |

**Figure S3.**
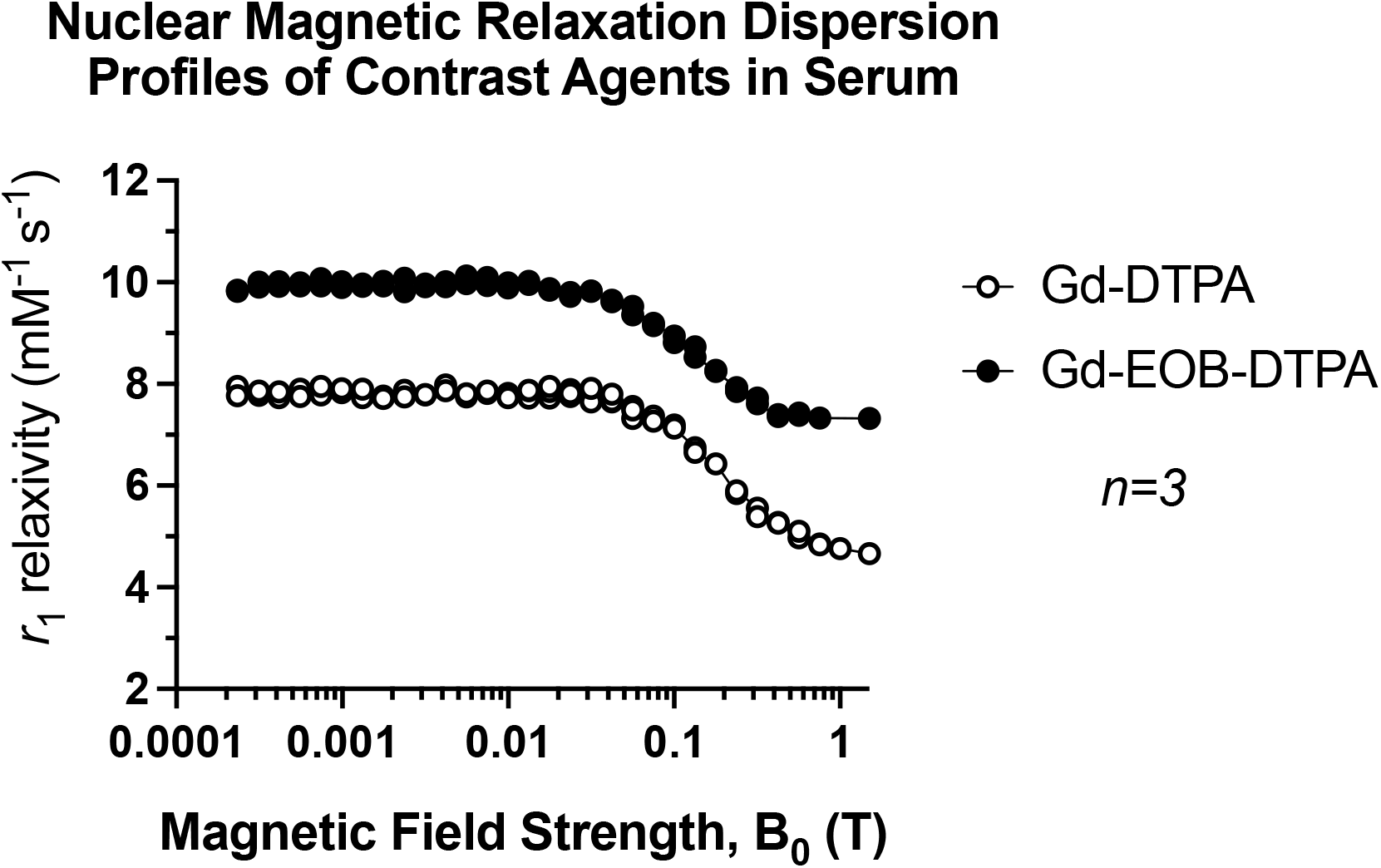
Nuclear magnetic relaxation dispersion (NMRD) profiles of Gd-DTPA and Gd-EOB-DTPA measured at 37°C. NMRD data for fields <1 T were acquired using a fast field-cycling relaxometer and extended by an inversion-recovery measurement at 1.5T on a clinical MRI system.

**Figure S4.**
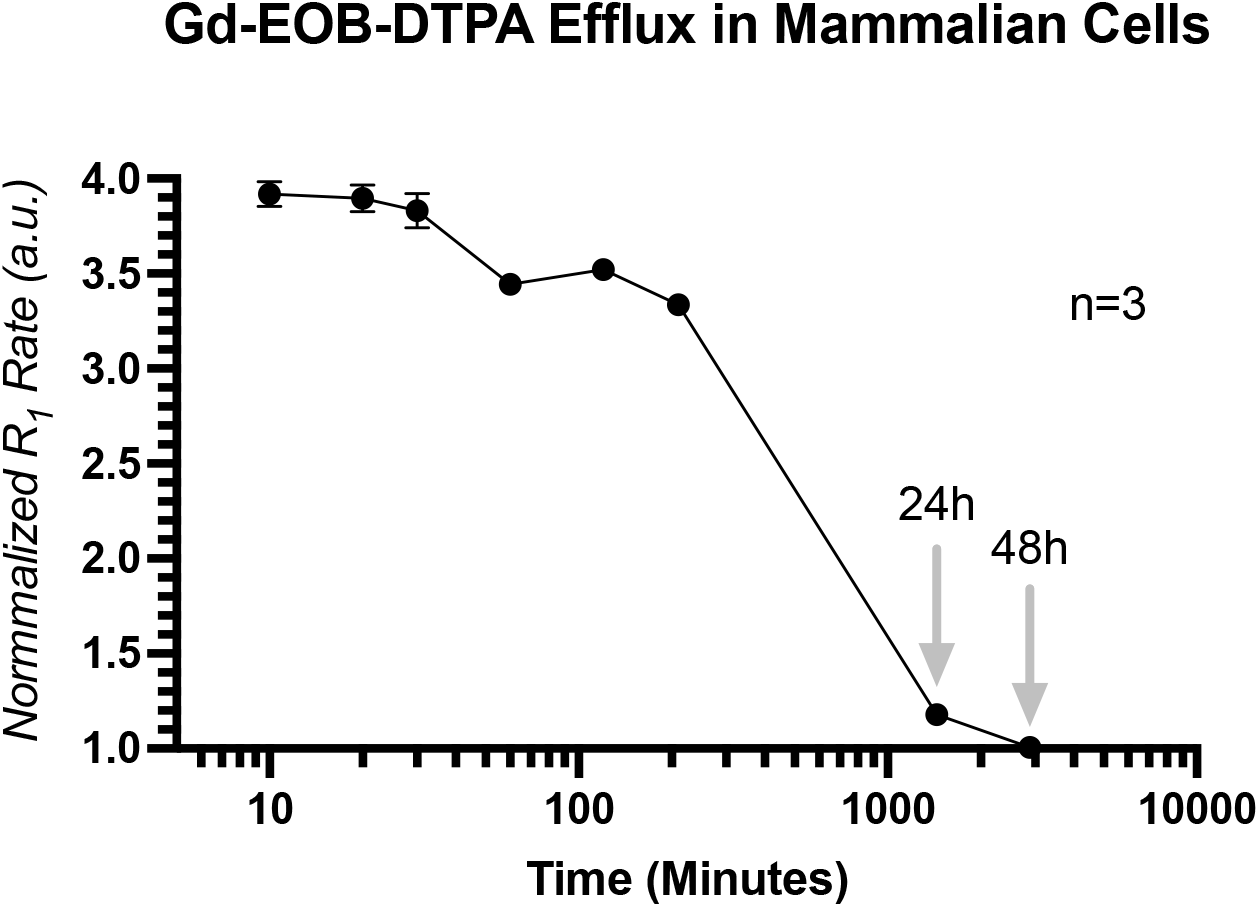
*R*_1_ relaxation rates of Luc-1B3 cells normalized to Luc-CTL cells (a.u.), measured at time, t following 90-minute incubation with 1.6 mM Gd-EOB-TPA and immediate wash with PBS. Measurements were acquired at 37°C, 10 minutes to 48 hours post-incubation. Error bars represent standard deviation. Some error bars are too small to represent on plot.

**Figure S5.**
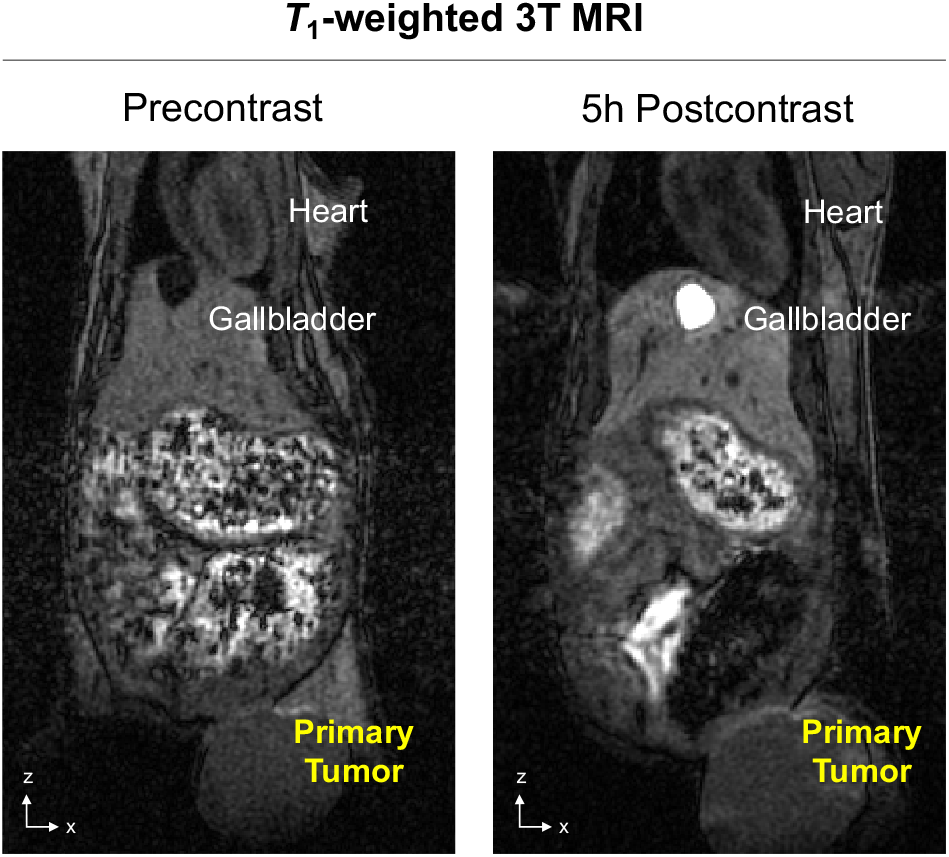
Pre-contrast and post-contrast (1.3 mmol/kg Gd-EOB-DTPA) *T*_1_-weighted images acquired at 3T of a representative Luc-CTL mouse on day 12.

**Figure S6.**
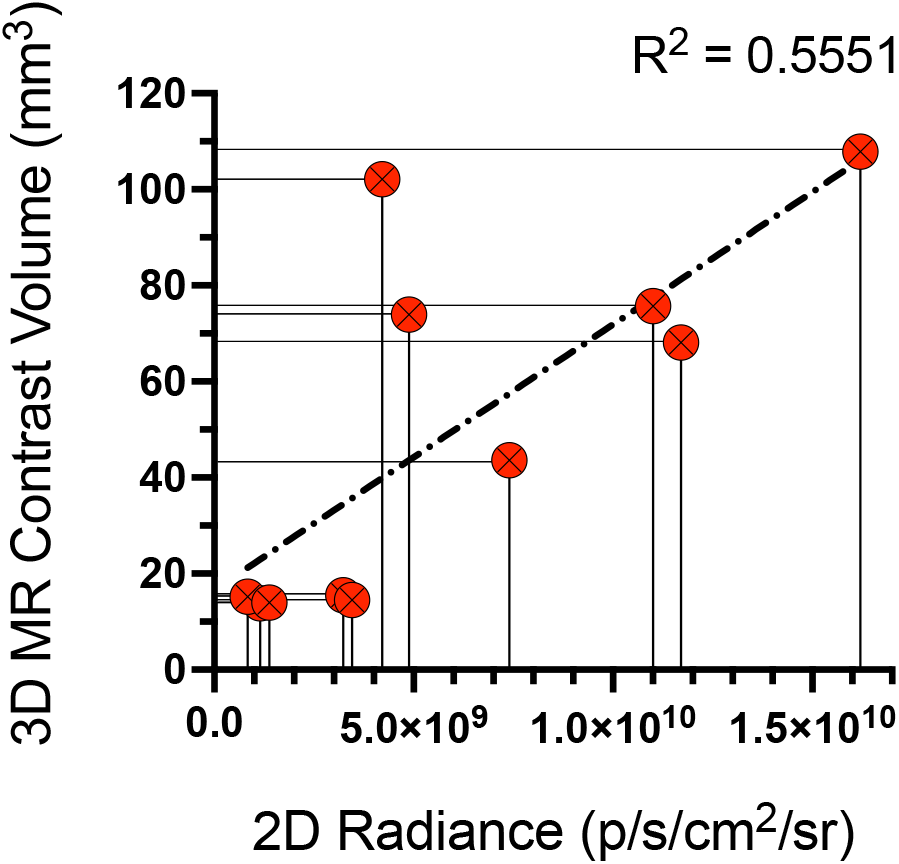
Correlation of 2D Radiance (p/s/cm^2^/sr) measured from primary tumors on Day 12 *versus* 3D MRI contrast-enhanced volume of the same primary tumors at the same timepoint.

**Figure S7.**
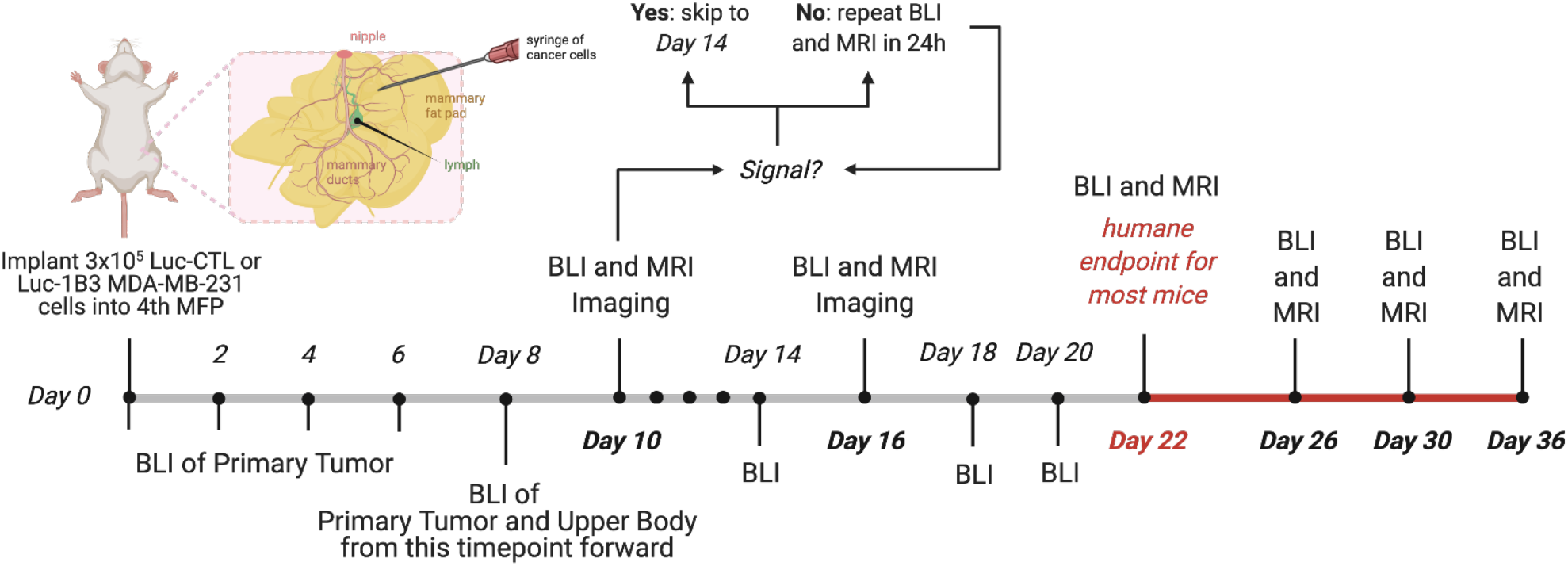
Timeline of metastasis and imaging protocol. Mammary fat pad, MFP. Bioluminescence imaging, BLI. *T*_1_-weighted magnetic resonance imaging at 3T, MRI.

**Figure S8.**
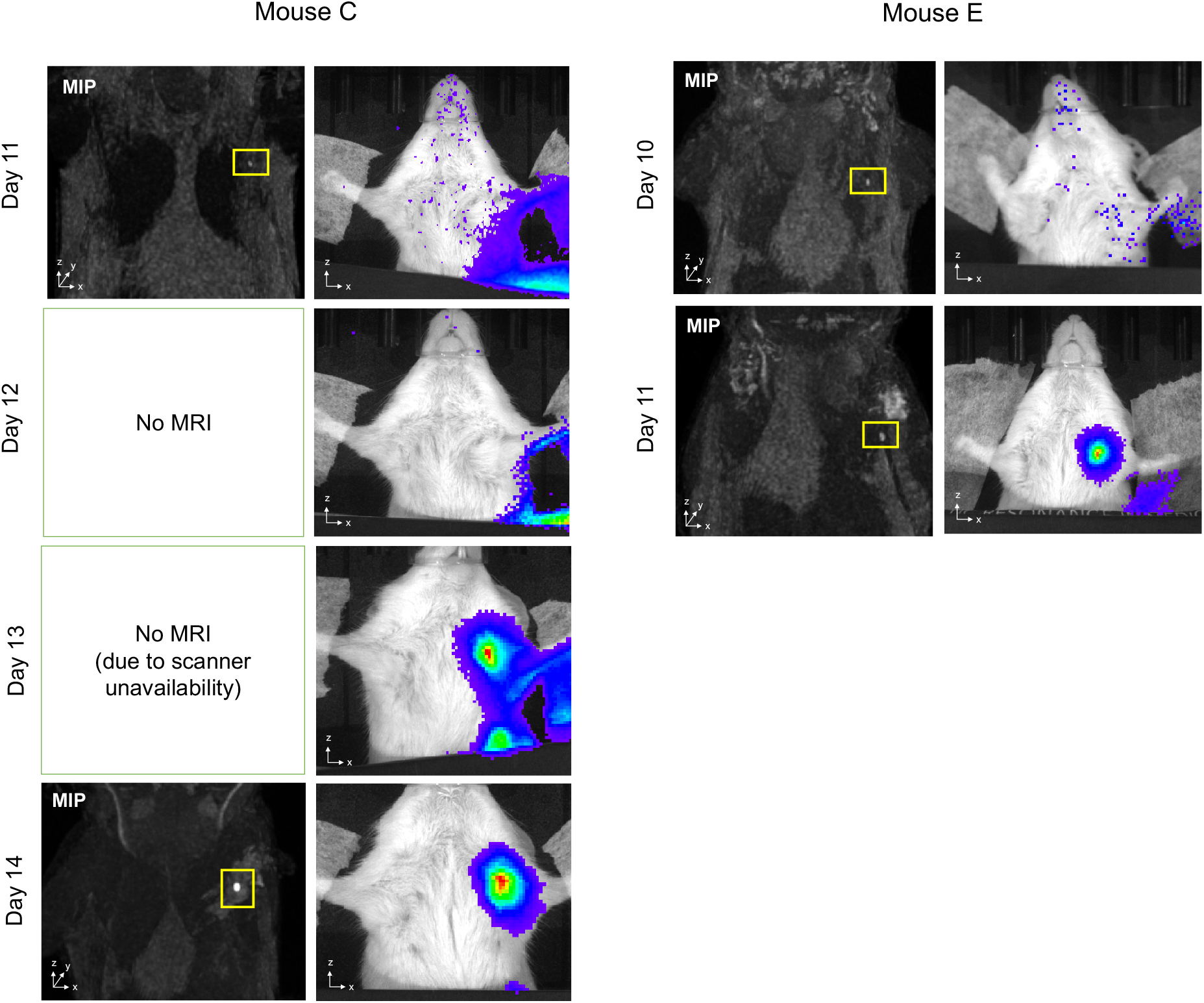
Bioluminescence images (BLI) and post-contrast (1.3 mmol/kg Gd-EOB-DTPA) *T*_1_-weighted images acquired at 3T of two mice, whereby detection of initial metastasis at the ipsilateral axillary lymph node was detected with MRI prior to detection with BLI.

## Notes

### Competing Interest Statement

The authors have declared no competing interest.

## REFERENCES

1 Lambert, A. W., Pattabiraman, D. R. & Weinberg, R. A. Emerging Biological Principles of Metastasis. Cell 168, 670–691, doi:10.1016/j.cell.2016.11.037 (2017).

2 Day, C. P., Merlino, G. & Van Dyke, T. Preclinical mouse cancer models: a maze of opportunities and challenges. Cell 163, 39–53, doi:10.1016/j.cell.2015.08.068 (2015).

3 Gengenbacher, N., Singhal, M. & Augustin, H. G. Preclinical mouse solid tumour models: status quo, challenges and perspectives. Nat Rev Cancer 17, 751–765, doi:10.1038/nrc.2017.92 (2017).

4 Francia, G., Cruz-Munoz, W., Man, S., Xu, P. & Kerbel, R. S. Mouse models of advanced spontaneous metastasis for experimental therapeutics. Nat Rev Cancer 11, 135–141, doi:10.1038/nrc3001 (2011).

5 Mezzanotte, L., van ‘t Root, M., Karatas, H., Goun, E. A. & Lowik, C. In Vivo Molecular Bioluminescence Imaging: New Tools and Applications. Trends Biotechnol 35, 640–652, doi:10.1016/j.tibtech.2017.03.012 (2017).

6 Giubellino, A. et al. Characterization of two mouse models of metastatic pheochromocytoma using bioluminescence imaging. Cancer Lett 316, 46–52, doi:10.1016/j.canlet.2011.10.019 (2012).

7 Taus, L. J., Flores, R. E. & Seyfried, T. N. Quantification of metastatic load in a syngeneic murine model of metastasis. Cancer Lett 405, 56–62, doi:10.1016/j.canlet.2017.07.011 (2017).

8 Kubota, S. I. et al. Whole-Body Profiling of Cancer Metastasis with Single-Cell Resolution. Cell Rep 20, 236–250, doi:10.1016/j.celrep.2017.06.010 (2017).

9 Pan, C. et al. Deep Learning Reveals Cancer Metastasis and Therapeutic Antibody Targeting in the Entire Body. Cell 179, 1661–1676 e1619, doi:10.1016/j.cell.2019.11.013 (2019).

10 Entenberg, D. et al. A permanent window for the murine lung enables high-resolution imaging of cancer metastasis. Nat Methods 15, 73–80, doi:10.1038/nmeth.4511 (2018).

11 Yan, C. et al. Visualizing Engrafted Human Cancer and Therapy Responses in Immunodeficient Zebrafish. Cell 177, 1903–1914 e1914, doi:10.1016/j.cell.2019.04.004 (2019).

12 Theruvath, J. et al. Anti-GD2 synergizes with CD47 blockade to mediate tumor eradication. Nat Med, doi:10.1038/s41591-021-01625-x (2022).

13 Anliang Xia, W. Y., Qiang Wang, Jianbo Xu, Yayun Gu, Liansheng Zhang, Chen Chen, Zhangding Wang, Di Wu, Qifeng He, Weiwei Yu, Fei Wang, Cailin Xue, Yan Zhang, Guojian Bao, Xuewen Tao, Siyuan Liu, Shouyu Wang, Zhibin Hu, Beicheng Sun. The cancer-testis lncRNA lnc-CTHCC promotes hepatocellular carcinogenesis by binding hnRNP K and activating YAP1 transcription. Nat Cancer, doi:https://doi.org/10.1038/s43018-021-00315-4 (2022).

14 Krupnick, A. S. et al. Quantitative monitoring of mouse lung tumors by magnetic resonance imaging. Nat Protoc 7, 128–142, doi:10.1038/nprot.2011.424 (2012).

15 Zhou, Z. et al. MRI detection of breast cancer micrometastases with a fibronectin-targeting contrast agent. Nat Commun 6, 7984, doi:10.1038/ncomms8984 (2015).

16 Louie, A. Y. et al. In vivo visualization of gene expression using magnetic resonance imaging. Nat Biotechnol 18, 321–325, doi:10.1038/73780 (2000).

17 Moore, A., Josephson, L., Bhorade, R. M., Basilion, J. P. & Weissleder, R. Human transferrin receptor gene as a marker gene for MR imaging. Radiology 221, 244–250, doi:10.1148/radiol.2211001784 (2001).

18 Cohen, B., Dafni, H., Meir, G., Harmelin, A. & Neeman, M. Ferritin as an endogenous MRI reporter for noninvasive imaging of gene expression in C6 glioma tumors. Neoplasia 7, 109–117, doi:10.1593/neo.04436 (2005).

19 Mukherjee, A., Wu, D., Davis, H. C. & Shapiro, M. G. Non-invasive imaging using reporter genes altering cellular water permeability. Nat Commun 7, 13891, doi:10.1038/ncomms13891 (2016).

20 Gilad, A. A. et al. Artificial reporter gene providing MRI contrast based on proton exchange. Nat Biotechnol 25, 217–219, doi:10.1038/nbt1277 (2007).

21 Shapiro, M. G. et al. Genetically encoded reporters for hyperpolarized xenon magnetic resonance imaging. Nat Chem 6, 629–634, doi:10.1038/nchem.1934 (2014).

22 Patrick, P. S. et al. Detection of transgene expression using hyperpolarized 13C urea and diffusion-weighted magnetic resonance spectroscopy. Magn Reson Med 73, 1401–1406, doi:10.1002/mrm.25254 (2015).

23 Patrick, P. S. et al. Dual-modality gene reporter for in vivo imaging. Proc Natl Acad Sci U S A 111, 415–420, doi:10.1073/pnas.1319000111 (2014).

24 Nystrom, N. N. et al. Longitudinal Visualization of Viable Cancer Cell Intratumoral Distribution in Mouse Models Using Oatp1a1-Enhanced Magnetic Resonance Imaging. Invest Radiol 54, 302–311, doi:10.1097/RLI.0000000000000542 (2019).

25 Kelly, J. J. et al. Safe harbor-targeted CRISPR-Cas9 homology-independent targeted integration for multimodality reporter gene-based cell tracking. Sci Adv 7, doi:10.1126/sciadv.abc3791 (2021).

26 Iwano, S. et al. Single-cell bioluminescence imaging of deep tissue in freely moving animals. Science 359, 935–939, doi:10.1126/science.aaq1067 (2018).

27 Sellmyer, M. A. et al. Imaging CAR T Cell Trafficking with eDHFR as a PET Reporter Gene. Mol Ther 28, 42–51, doi:10.1016/j.ymthe.2019.10.007 (2020).

28 Chapy, H. et al. PBPK modeling of irbesartan: incorporation of hepatic uptake. Biopharm Drug Dispos 36, 491–506, doi:10.1002/bdd.1961 (2015).

29 Powell, J. et al. Novel mechanism of impaired function of organic anion-transporting polypeptide 1B3 in human hepatocytes: post-translational regulation of OATP1B3 by protein kinase C activation. Drug Metab Dispos 42, 1964–1970, doi:10.1124/dmd.114.056945 (2014).

30 Gasteiger, E. et al. ExPASy: The proteomics server for in-depth protein knowledge and analysis. Nucleic Acids Res 31, 3784–3788, doi:10.1093/nar/gkg563 (2003).

31 Alam, K. et al. Regulation of Organic Anion Transporting Polypeptides (OATP) 1B1-and OATP1B3-Mediated Transport: An Updated Review in the Context of OATP-Mediated Drug-Drug Interactions. Int J Mol Sci 19, doi:10.3390/ijms19030855 (2018).

32 Jumper, J. et al. Highly accurate protein structure prediction with AlphaFold. Nature 596, 583–589, doi:10.1038/s41586-021-03819-2 (2021).

33 Huang, H. et al. iPTMnet: an integrated resource for protein post-translational modification network discovery. Nucleic Acids Res 46, D542–D550, doi:10.1093/nar/gkx1104 (2018).

34 York, W. S. et al. GlyGen: Computational and Informatics Resources for Glycoscience. Glycobiology 30, 72–73, doi:10.1093/glycob/cwz080 (2020).

35 Hornbeck, P. V. et al. PhosphoSitePlus: a comprehensive resource for investigating the structure and function of experimentally determined post-translational modifications in man and mouse. Nucleic Acids Res 40, D261–270, doi:10.1093/nar/gkr1122 (2012).

36 Miraux, S., Massot, P., Ribot, E. J., Franconi, J. M. & Thiaudiere, E. 3D TrueFISP imaging of mouse brain at 4.7T and 9.4T. J Magn Reson Imaging 28, 497–503, doi:10.1002/jmri.21449 (2008).

37 Szomolanyi, P. et al. Comparison of the Relaxivities of Macrocyclic Gadolinium-Based Contrast Agents in Human Plasma at 1.5, 3, and 7 T, and Blood at 3 T. Invest Radiol 54, 559–564, doi:10.1097/RLI.0000000000000577 (2019).

38 Puchalapalli, M. et al. NSG Mice Provide a Better Spontaneous Model of Breast Cancer Metastasis than Athymic (Nude) Mice. PLoS One 11, e0163521, doi:10.1371/journal.pone.0163521 (2016).

39 Del Monte, U. Does the cell number 10(9) still really fit one gram of tumor tissue? Cell Cycle 8, 505–506, doi:10.4161/cc.8.3.7608 (2009).

40 Deoni, S. C., Peters, T. M. & Rutt, B. K. High-resolution T1 and T2 mapping of the brain in a clinically acceptable time with DESPOT1 and DESPOT2. Magn Reson Med 53, 237–241, doi:10.1002/mrm.20314 (2005).

41 Luetkens, J. A. et al. Quantification of Liver Fibrosis at T1 and T2 Mapping with Extracellular Volume Fraction MRI: Preclinical Results. Radiology 288, 748–754, doi:10.1148/radiol.2018180051 (2018).

42 Richardson, D. S. & Lichtman, J. W. Clarifying Tissue Clearing. Cell 162, 246–257, doi:10.1016/j.cell.2015.06.067 (2015).

43 Contag, C. H., Jenkins, D., Contag, P. R. & Negrin, R. S. Use of reporter genes for optical measurements of neoplastic disease in vivo. Neoplasia 2, 41–52, doi:10.1038/sj.neo.7900079 (2000).

44 Hung, K. E. et al. Development of a mouse model for sporadic and metastatic colon tumors and its use in assessing drug treatment. Proc Natl Acad Sci U S A 107, 1565–1570, doi:10.1073/pnas.0908682107 (2010).

45 Anderson, R. L. et al. A framework for the development of effective anti-metastatic agents. Nat Rev Clin Oncol 16, 185–204, doi:10.1038/s41571-018-0134-8 (2019).

46 Martin, O. A., Anderson, R. L., Narayan, K. & MacManus, M. P. Does the mobilization of circulating tumour cells during cancer therapy cause metastasis? Nat Rev Clin Oncol 14, 32–44, doi:10.1038/nrclinonc.2016.128 (2017).

47 Obradovic, M. M. S. et al. Glucocorticoids promote breast cancer metastasis. Nature 567, 540–544, doi:10.1038/s41586-019-1019-4 (2019).

48 Obenauf, A. C. et al. Therapy-induced tumour secretomes promote resistance and tumour progression. Nature 520, 368–372, doi:10.1038/nature14336 (2015).

49 Tanaka, Y. et al. Sentinel Lymph Node-Targeted Therapy by Oncolytic Sendai Virus Suppresses Micrometastasis of Head and Neck Squamous Cell Carcinoma in an Orthotopic Nude Mouse Model. Mol Cancer Ther 18, 1430–1438, doi:10.1158/1535-7163.MCT-18-1372 (2019).

50 Esposito, M., Ganesan, S. & Kang, Y. Emerging strategies for treating metastasis. Nat Cancer 2, 258–270, doi:10.1038/s43018-021-00181-0 (2021).

51 Meirson, T., Gil-Henn, H. & Samson, A. O. Invasion and metastasis: the elusive hallmark of cancer. Oncogene 39, 2024–2026, doi:10.1038/s41388-019-1110-1 (2020).

52 Kluger, H. M. et al. Using a xenograft model of human breast cancer metastasis to find genes associated with clinically aggressive disease. Cancer Res 65, 5578–5587, doi:10.1158/0008-5472.CAN-05-0108 (2005).

53 Kiryu, S. et al. Diet and gastrointestinal signal on T1-weighted magnetic resonance imaging of mice. Magn Reson Imaging 28, 273–280, doi:10.1016/j.mri.2009.10.005 (2010).

54 Nivin N. Nyström, H. L., Francisco M. Martinez, Xiao-an Zhang, Timothy J. Scholl, John A. Ronald. Gadolinium-free Magnetic Resonance Imaging of the Liver using an Oatp1-targeted Manganese(III) Porphyrin. bioRxiv 2021.08.04.455144, doi:https://doi.org/10.1101/2021.08.04.455144 (2021).

55 Arvanitis, C. D., Ferraro, G. B. & Jain, R. K. The blood-brain barrier and blood-tumour barrier in brain tumours and metastases. Nat Rev Cancer 20, 26–41, doi:10.1038/s41568-019-0205-x (2020).

56 Honkala, A., Malhotra, S. V., Kummar, S. & Junttila, M. R. Harnessing the predictive power of preclinical models for oncology drug development. Nat Rev Drug Discov 21, 99–114, doi:10.1038/s41573-021-00301-6 (2022).

57 Nivin N. Nyström, L. C. M. Y., Jeffrey J.L. Carson, Timothy J. Scholl, John A. Ronald Development of a Human Photoacoustic Imaging Reporter Gene Using the Clinical Dye Indocyanine Green. Radiology: Imaging Cancer 1, e190035, doi:https://doi.org/10.1148/rycan.2019190035 (2019).

58 Rydberg, J. N. et al. T1-weighted MR imaging of the brain using a fast inversion recovery pulse sequence. J Magn Reson Imaging 6, 356–362, doi:10.1002/jmri.1880060216 (1996).

59 Bydder, G. M. & Young, I. R. MR imaging: clinical use of the inversion recovery sequence. J Comput Assist Tomogr 9, 659–675 (1985).

60 Yushkevich, P. A. et al. User-guided 3D active contour segmentation of anatomical structures: significantly improved efficiency and reliability. Neuroimage 31, 1116–1128, doi:10.1016/j.neuroimage.2006.01.015 (2006).

